# Defective flow-migration coupling causes arteriovenous malformations in hereditary hemorrhagic telangiectasia

**DOI:** 10.1101/2021.05.06.442985

**Authors:** Hyojin Park, Jessica Furtado, Mathilde Poulet, Minhwan Chung, Sanguk Yun, Sungwoon Lee, William C Sessa, Claudio Franco, Martin A Schwartz, Anne Eichmann

## Abstract

**Background:** Activin receptor-like kinase 1 (*ACVRL1*, hereafter *ALK1*) is an endothelial transmembrane serine threonine kinase receptor for BMP family ligands that plays a critical role in cardiovascular development and pathology. Loss-of-function mutations in the *ALK1* gene cause type 2 hereditary hemorrhagic telangiectasia (HHT), a devastating disorder that leads to arteriovenous malformations (AVMs). Here we show that ALK1 controls endothelial cell polarization against the direction of blood flow and flow-induced endothelial migration from veins through capillaries into arterioles.

**Methods:** Using Cre lines that recombine in different subsets of arterial, capillary-venous or endothelial tip cells, we showed that capillary-venous *Alk1* deletion was sufficient to induce AVM formation in the postnatal retina.

**Results:** ALK1 deletion impaired capillary-venous endothelial cell polarization against the direction of blood flow *in vivo* and *in vitro*. Mechanistically, ALK1 deficient cells exhibited increased integrin signaling interaction with VEGFR2, which enhanced downstream YAP/TAZ nuclear translocation. Pharmacological inhibition of integrin or YAP/TAZ signaling rescued flow migration coupling and prevented vascular malformations in *Alk1* deficient mice.

**Conclusions:** Our study reveals ALK1 as an essential driver of flow-induced endothelial cell migration and identifies loss of flow-migration coupling as a driver of AVM formation in HHT disease. Integrin-YAP/TAZ signaling blockers are new potential targets to prevent vascular malformations in HHT patients.

## Introduction

Hereditary hemorrhagic telangiectasia (HHT) is an inherited autosomal dominant vascular disorder that causes arteriovenous malformations (AVMs) in more than 1.4 million people worldwide^1^. More than 80 % of HHT cases are caused by heterozygous mutations in the endothelial surface receptors Endoglin (*ENG,* mutated in HHT1) and *ACVRL1* (hereafter referred to as *ALK1*, mutated in HHT2), and mutations in *SMAD4* cause a combined juvenile polyposis-HHT syndrome that accounts for <5% of HHT cases^2–5^. ALK1 and ENG are receptors for TGF-β superfamily members BMP9 and BMP 10^6, 7^. Ligand binding activates ALK1/ENG receptor signaling to cytoplasmic SMAD 1/5/8, which subsequently complex with SMAD4 and translocate into the nucleus to regulate gene expression^8^. Thus, known HHT mutations affect different components of an endothelial signaling pathway that prevents vessels from forming AVMs. A recent study has shown that somatic second-hits inactivating the remaining intact *ALK1* or *ENG* allele occurred in the lesions, supporting that vascular malformations in HHT are caused by a two-hit mechanism^9^.

Whereas the genetics of AVM have been well studied, the underlying cellular and molecular principles are not fully understood, thus limiting the development of new treatment options. AVMs are direct connections between arteries and veins that lack an intermediate capillary bed^2^. AVMs in HHT patients appear most often in the skin, oral cavity, nasal, and gastrointestinal (GI) tract mucosa, lung, liver, and brain. Small AVMs in the skin and mucus membranes are called telangiectasias; rupture of these lesions leads to frequent epistaxis, GI bleeding, and anemia, all of which are major quality of life issues for HHT patients^10^. Larger AVMs in liver, lung, or brain may additionally cause life-threatening conditions such as high output heart failure and stroke^11^. We and others previously showed that pan-endothelial knockout of *Alk1* using *Alk1^f/f^ Cdh5 Cre^ERT2^* in neonates led to AVMs in retina, brain and internal organs, indicating that endothelial ALK1 is necessary for proper vascular development^12, 13^. However, what types of ECs are responsible and how AVMs develop remains largely unknown.

Previous data from us and others have shown that BMP9/10-ALK1-ENG-SMAD4 signaling is enhanced by flow, and initiates a negative feedback signal that dampens flow-induced activation of AKT, thereby coordinating proper vascular remodeling^12, 14–17^. Mechanistically, blocking BMP9-ALK1-ENG signaling promotes endothelial PI3K (phosphatidylinositol 3-kinase)/AKT activation. ALK1-deficient ECs showed enhanced phosphorylation of the PI3K target AKT and vascular endothelial growth factor receptor 2 (VEGFR2)^13, 18^. Pharmacological VEGFR2 or PI3K inhibition prevented AVM formation in *Alk1*-deficient mice and decreased diameter of AVMs in ENG mutants^19^. Moreover, an increase in PI3K signaling has been recently confirmed in cutaneous telangiectasia biopsies of patients with HHT2^20, 21^.

Here we investigated the origin of AVM-causing cells using novel Cre lines that delete *Alk1* in subsets of ECs. In doing so, we observed that ECs in remodeling vessels move against the direction of blood flow, while maintaining vascular integrity. In response to the physical forces such as wall shear stress exerted by blood, ECs polarize their Golgi apparatus in front of the nucleus (front-rear polarity) and migrate against the blood flow from veins towards arteries. We further provide evidence that ALK1 contributes to flow-migration coupling via VEGFR2-integrin signaling and downstream YAP/TAZ nuclear translocation. Collectively, the data show that ALK1 controls flow-induced cell migration to prevent AVM formation and identify new targets with the potential to prevent vascular malformations in HHT patients.

### Methods Mice

All animal experiments were performed under a protocol approved by Institutional Animal Care Use Committee of Yale University. *Alk1^f/f^* mice were kindly provided by Dr. S. Paul Oh. *Bmx Cre^ERT2^* and *Esm1 Cre^ERT2^* mice were kindly provided by Dr. Ralf Adams and *Mfsd2a Cre^ERT2^* mice were kindly provided by Dr. Bin Zhou. Seven to eight weeks old *Alk1^f/f^* and *Bmx Cre^ERT2^ mTmG*, *Esm1 Cre^ERT2^ mTmG* mice or *Mfsd2a Cre^ERT2^ mTmG* mixed genetic background were intercrossed for experiments and *Alk1^f/f^ Mfsd2a Cre^ERT2^ (mTmG)*, *Alk1^f/f^ Esm1 Cre^ERT2^ (mTmG)* mice or *Alk1^f/f^ Bmx Cre^ERT2^ (mTmG)* were used. Gene deletion was induced by intra-gastric injections with 100 μg Tx (Sigma, T5648; 2.5 mg ml^−1^) into pups at P4 or P1-3. Tx-injected *Cre^ERT2^* negative littermates were used as controls.

### Latex dye injection

P6 pups were anaesthetized on ice, and abdominal and thoracic cavities were opened. The right atrium was cut, blood was washed out with 2 ml PBS and 1 ml of latex dye was slowly and steadily injected into the left ventricle with an insulin syringe. Retinas and GI tracts were washed in PBS and fixed with 4% paraformaldehyde (PFA) overnight. Brains and GI tracts were cleared in Benzyl Alcohol: Benzyl Bezonate (1:1) for 2-3 days before imaging.

### Reagents and antibodies

For immunostaining: IB4 ([IsolectinB4] #121412, 10 μg/mL; Life Technologies), GFP Polyclonal Antibody, Alexa Fluor 488 (#A-21311, 1:1000; Invitrogen), GOLPH4 (#ab28049, 1:400; abcam), anti-YAP (#14074, 1:300; Cell Signaling), anti-TAZ (#HPA007415, 1:300 Sigma), mouse anti-ALK1 (#AF770, 1:300; R&D) human anti-ALK1(#AF370, 1:300; R&D), VE-Cadherin (#555289, 1:200; BD) GM-130 (#610822, 1:500; BD), DAPI (#D1306, 1:1000; Life Technologies), anti-integrin β1 Alexa Fluor 647 (#303047,1:500; BioLegend), anti-integrin α5 and αv (From Martin A Schwartz) For western blotting: anti-ALK1 (7R-49334, 1:1000; Fitzgerald), anti-integrin β1 (#34971, Cell Signaling), anti-integrin α5 and αv, anti-VEGFR2 (#9698, Cell Signaling), β-actin (#A1978 1:3000; Sigma), anti-YAP (#14074, 1:1000; Cell Signaling), anti-TAZ (#HPA007415, 1:2000 Sigma).

Appropriate secondary antibodies were fluorescently labeled (Alexa Fluor donkey anti-rabbit, Alexa Fluor donkey anti-goat) or conjugated to horseradish peroxidase (anti-rabbit and anti-mouse IgG [H+L], 1:8.000; Vector Laboratories). ATN-161 (#S8454, Selleckchem), cilengitide trifluoroacetate (#S7077, Selleckchem), verteporfin (#S1786, Selleckchem), wortmannin (#S2758, Selleckchem)

### Immunostaining

For angiogenesis studies the eyes of P6/P8 pups were prefixed in 4% PFA for 8 min at room temperature. Retinas were dissected, blocked for 30 min at room temperature in blocking buffer (1% fetal bovine serum, 3% BSA, 0.5% Triton X-100, 0.01% Na deoxycholate, 0.02% Sodium Azide in PBS at pH 7.4) and then incubated with specific antibodies in blocking buffer overnight at 4°C. The next day, retinas were washed and incubated with IB4 together with the corresponding secondary antibody for overnight at 4°C. The next day, retinas were washed and post-fixed with 0.1% PFA and mounted in fluorescent mounting medium (DAKO, USA). High-resolution pictures were acquired using ZEISS LSM800 and Leica SP8 confocal microscope with a Leica spectral detection system (Leica TCS SP8 detector), and the Leica application suite advanced fluorescence software. Quantification of retinal vasculature was done using ImageJ and then Prism 7 software for statistical analysis.

For cell immunostaining, cells were plated on gelatin coated dishes. Growing cells were fixed for 10 min with 4% paraformaldehyde (PFA) and permeabilized with 0.1% Triton X-100 for 10 min prior to overnight incubation with primary antibody and then secondary antibody conjugated with fluorophore.

### Cell culture and siRNA transfection

Human umbilical vein endothelial cells (HUVECs) were obtained from the Yale University Vascular Biology and Therapeutics Core Facility and cultured in EGM2-Bullet kit medium (CC-3156 & CC-4176, Lonza). Depletion of *ALK1*, *SMAD4* or *ENG* was achieved by transfecting 20 pmol of small interfering RNA (siRNA) against *ALK1* (Qiagen, mixture of 2 siRNAs: S102659972 and S102758392), *SMAD4* (Dharmacon, SMARTpool: ON-TARGETplus L-003902-00-0005) or *ENG* (Dharmacon, ON-TARGETplus LQ-011026-00-0005) using Lipofectamine RNAiMax (Invitrogen). Transfection efficiency was assessed by western blotting and quantitative PCR (qPCR). Experiments were performed 60 hours posttransfection and results were compared with siRNA CTRL (ON-TARGETplus Non-Targeting Pool D-001810-10-05).

### Shear stress experiments

HUVECs were re-plated on glass slides coated with the indicated proteins for 6 hours and wound scratch was carried out on the slides before application of flow. The slides were loaded into parallel plate flow chambers. Laminar shear at 15 dynes/cm^2^ was used to mimic high flow in retinal veins.

For real-time imaging, PH-AKT-mClover3 was modified from PH-AKT-GFP (addgene #51465). Plasma membrane targeting sequence of LCK tagged with mRuby3 (LCK-mRuby3, modified from addgene #98822) was co-expressed in the same vector by IRES sequence as a plasma membrane marker. HUVECs were transfected with siRNAs followed by lentiviral transduction coding PH-AKT-mClover3 and LCK-mRuby3. The infected cells were mixed with uninfected cells in 1:2 ratio, then seeded on microfluidic chamber (IBIDI u-slide 0.4 luer, 1×10^5^ total cell/slide) and cultured additional 24-48 hours more for imaging. Imaging was performed on an Eclipse Ti microscope equipped with an Ultraview Vox spinning disk confocal imaging system, with 20X objective (Plan Apo, Nikon). Each pixel intensity was plotted as y with corresponding distance from upstream of the cell as x, which was normalized to have length 1 to the direction of flow (0 to 1). Then slope of the plot at each time frame was measured as a representative value of cell polarity.

### Western blotting

Cells were lysed with Laemmli buffer including phosphatase and protease inhibitors (Thermo Scientific, 78420, 1862209). 20 μg of proteins were separated on 4% to 15% Criterion precast gels (567–1084, Biorad) and transferred on 0.23 um nitrocellulose membranes (Biorad). Western blots were developed with chemiluminescence horseradish peroxidase substrate (Millipore, WBKLS0500) on a Luminescent image Analyzer, ImageQuant LAS 4000 mini (GE Healthcare). Bands were quantified using ImageJ.

### Immunoprecipitation

Cell lysates were prepared in 50 mM Tris-HCl at pH 7.4, 50 mM NaCl, 0.5% Triton X-100, phosphatase and protease inhibitors, centrifuged at 16,000 × *g* for 20 min. Protein concentration was quantified using Bradford assay (Pierce). In total, 500 μg of protein from cell lysate were incubated overnight at 4 °C with 10 μg/ml of anti-VEGFR2, and finally incubated with protein A/G magnetic beads (88802, Thermo Scientific) for 2 h at 4 °C. The immunocomplexes were washed three times in lysis buffer and resuspended in 1X Laemmli’s sample buffer. For western-blot analysis, 50 μg of protein was loaded for each condition.

### Polarity index calculation

Briefly, after segmenting each channel corresponding to the Golgi and nuclear staining, the centroid of each organelle was determined and a vector connecting the center of the nucleus to the center of its corresponding Golgi apparatus was drawn. The Golgi-nucleus assignment was done automatically minimizing the distance between all the possible couples. The polarity of each cell was defined as the angle between the vector and the scratch line. An angular histogram showing the angle distribution was then generated. Circular statistic was performed using the Circular Statistic Toolbox. To test for circular uniformity, we applied the polarity index (PI), calculated as the length of mean resultant vector for a given angular distribution^22^.

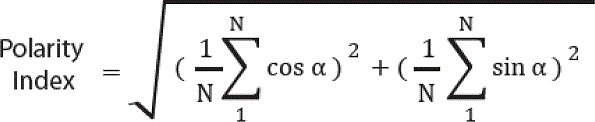

### Statistical Analysis

All data are shown as mean standard error of the mean. Tow-tailed unpaired t-test was used to compare 2 groups. Oneway ANOVA was used to compare more than 2 groups followed by appropriate post hoc multiple comparison procedure (Holm-Sidak multiple comparisons test). To construct the survival curves we have used the Kaplan–Meier method. P value <0.05 was considered to be statistically significant. Statistical analyses were performed for all quantitative data using Prism 6.0 (Graph Pad).

## Results

### Endothelial lineage tracing reveals flow migration coupling in retinal vessels

To track the dynamics of endothelial flow migration coupling in the mouse retina, we used three *Cre^ERT2^* lines that recombine in subsets of ECs. These include Major Facilitator Superfamily Domain Containing 2a (*Mfsd2a*), which recombines venous and capillary ECs but not arteries or tip cells in the brain vasculature^23–25^; Endothelial cell-specific molecule 1 (*Esm1*), which recombines tip cells and their progeny^26, 27^; and the artery-specific Bone marrow x (*Bmx*)^28^ line. We intercrossed these lines with *mTmG* reporter mice^29^ to lineage-trace GFP positive *Mfsd2a*, *Esm1* and *Bmx* expressing ECs. Tamoxifen (Tx) was injected 12 h, 24 h and 48 h prior to sacrifice at P6 (Figure 1 A-I). At 12 h post injection, *Mfsd2a* positive cells were absent from the tip cell position, but labeled veins and capillaries in the vascular plexus, as well as the distal pole of arterioles (Figure 1A). *Esm1*-positive cells were restricted to the tip position, while *Bmx Cre^ERT2^* positive cells were located in the proximal part of retinal arterioles close to the optic nerve (Figure 1B-C). Hence the three *Cre* lines labeled distinct and non-overlapping endothelial cell populations at this time point.

**Figure 1.**
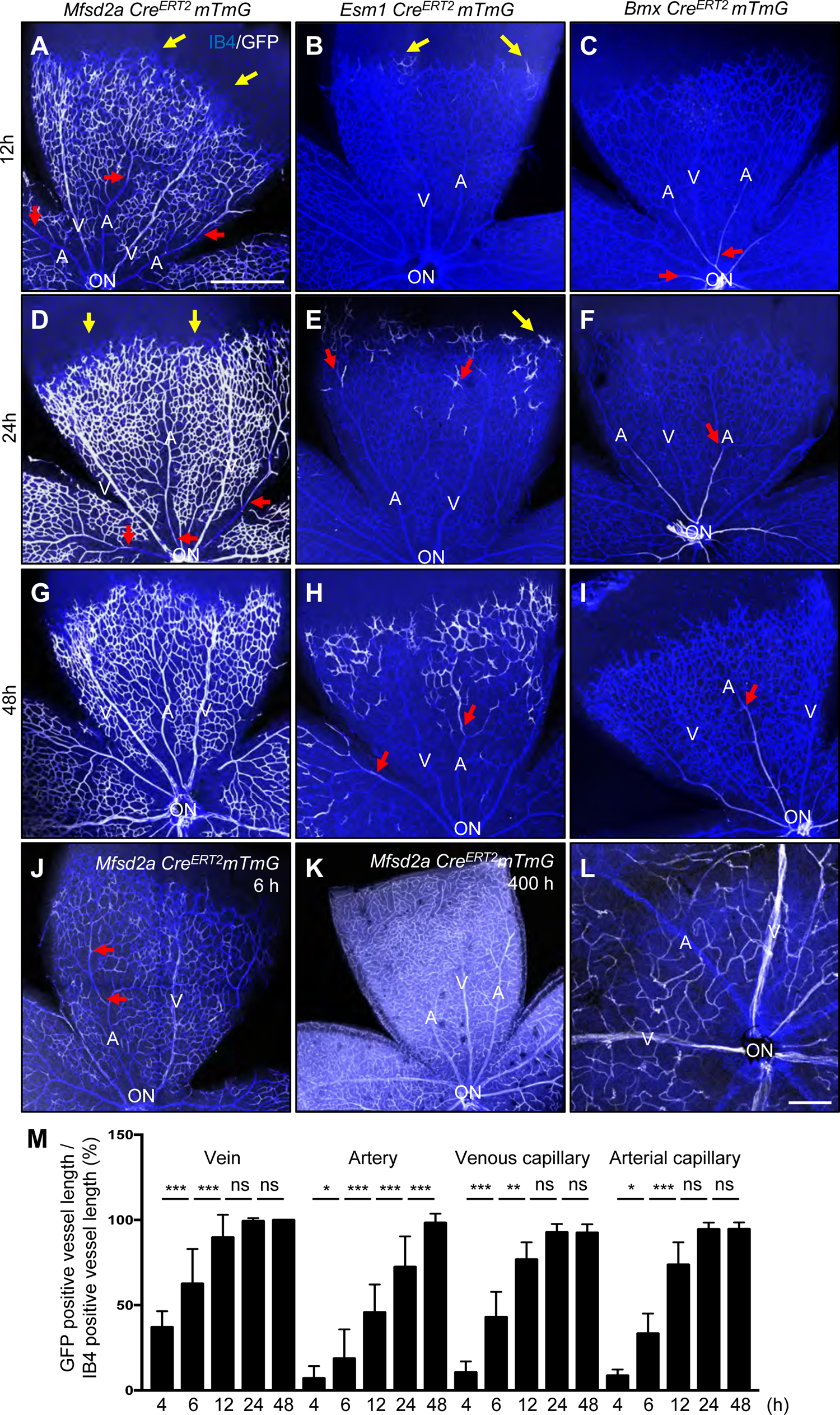
Retinal endothelial cell lineage tracing. (A-I) P6 retina flat mount images labeled with IB4 (blue) and GFP (white) from *Mfsd2a Cre^ERT2^ mTmG* (A, D, G), *Esm1 Cre^ERT2^ mTmG* (B, E, H) and *Bmx Cre^ERT2^ mTmG* (C, F, I) mice injected with 100 μg Tx at P5.5 (12 h, A-C), P5 (for 24 h, D-F) and P4 (for 48 h, G-I) and dissected at P6. (J) 100 μg Tx was injected at P6 and dissected after 6 h in *Mfsd2a Cre^ERT2^ mTmG* mice. (K) 100 μg Tx was injected at P4 and dissected after 400 h (P21) (L) 2 mg/kg Tx was injected in P20 *Mfsd2a Cre^ERT2^ mTmG* mice and dissect at P21. Yellow arrows indicate tip cells and red arrows indicate location of GFP-expressing ECs in arteries. (M) Quantification of *Mfsd2a Cre^ERT2^ mTmG* GFP expressing vessel length over IB4 positive vessel length from optic nerve. n = 6-8 retinas per time point. P-value < 0.001, Error bars: SEM. *P-value < 0.05, **P-value < 0.01, ***P-value < 0.001, ns: nonsignificant, One-way ANOVA. ON: optic nerve, V: vein, A: artery, Scale bars: 500 μm (A-K) and 50 μm (L).

24 h and 48 h after injection, *Mfsd2a*-positive cells were still excluded from the tip position, but progressively colonized the arteries from the distal to the proximal part (Figure 1D,G). *Esm1*-positive cells were seen at the tip position and moving towards the distal parts of the arterioles at 24h and 48 h after injection (Figure 1E,H), while *Bmx*-positive cells remained confined to the proximal arterioles (Figure 1F,I). Very few *Mfsd2a*-GFP positive cells were detected in arteries 4 h and 6 h post Tx injection (Supp. Figure 1 A, Figure 1J), while 400 h after P4 Tx injection, ie at P21, most of the retinal endothelium was GFP positive (Figure 1K). By contrast, a single Tx injection at P20 labeled venous and capillary endothelium, but not arteries at P21 (Figure 1 L), demonstrating that ECs of venous and capillary origin migrate against the direction of flow into neighboring arteries during vascular remodeling.

To quantify displacement of *Msfd2a*-GFP positive cells, we measured the relative length of GFP positive area in retinal arteries, veins and capillaries at different time points (Figure 1M). After 12 h, about 90 % of venous and capillary vessel area was occupied by GFP-positive cells, while only 50% of the distal arterial vessel area was occupied by GFP-positive cells and this gradually increased over time until 48 h (Figure 1M), demonstrating quantifiable displacement of capillary and venous ECs towards arteries over time.

### *Alk1* deletion in capillary and venous ECs causes AVMs

To determine the origin of AVM forming cells in *Alk1* mutants, we next intercrossed *Mfsd2a*, *Esm1* and *Bmx Cre^ERT2^* mice with *Alk1^f/f^ mTmG* reporter mice. Tx was injected at P4 and mice were analyzed at P6 (Figure 2 A). Efficient Alk1 deletion was verified in all three lines using immunostaining (Supp. Figure 2). Interestingly, venous and capillary endothelial *Alk1* deletion using the *Mfsd2a Cre^ERT2^* driver line led to numerous AVMs in the retina (Figure 2 B and E). By contrast, neither *Alk1^f/f^ Esm1 Cre^ERT2^* nor *Alk1^f/f^ Bmx Cre^ERT2^* mutants displayed any retinal AVMs (Figure 2 C-D and F-G). We analyzed the presence of retinal and brain AVMs by injection of latex dye into the left ventricle of P6 *Alk1^f/f^ Mfsd2a Cre^ERT2^* and control littermates (Figure 2 H-K). The latex dye does not cross the capillary beds and was retained within the arterial branches in *Alk1^f/f^* brain and retina (Figure 2 H-I). In the *Alk1^f/f^ Mfsd2a Cre^ERT2^* mutants, the latex penetrated both the venous as well as the arterial branches via AVMs in the retina and brain (Figure 2 J-K). To see whether *Alk1^f/f^ Esm1 Cre^ERT2^* and *Alk1^f/f^ Bmx Cre^ERT2^* could develop AVMs by longer-term exposure of Tx, Tx was injected at P1 and mice were analyzed at P6 (Figure 2 L). Neither *Alk1^f/f^ Esm1 Cre^ERT2^* nor *Alk1^f/f^ Bmx Cre^ERT2^* mutants exhibited any AVMs (Figure 2 M-N).

**Figure 2.**
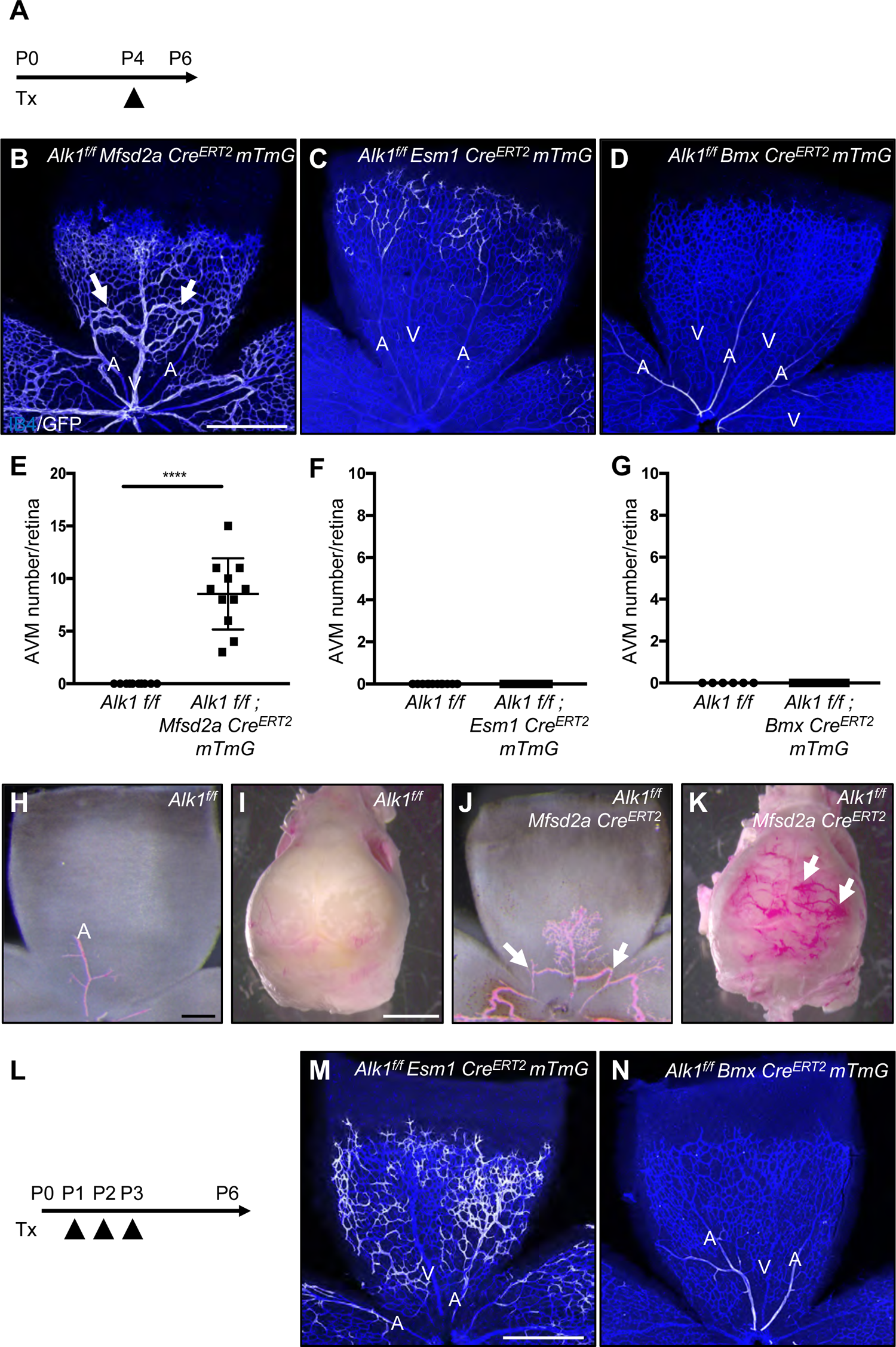
Capillary-venous loss of ALK1 leads to retinal and brain AVMs. (A) Schematic representation of the experimental strategy used to delete *Alk1* in mice (P4-P6). (B-D) P6 retina flat mount images labeled with IB4 (blue) and GFP (white) from *Alk1^f/f^ Mfsd2a Cre^ERT2^ mTmG* (B), *Alk1^f/f^ Esm1 Cre^ERT2^ mTmG* (C) and *Alk1^f/f^ Bmx Cre^ERT2^ mTmG* pups (D) injected with 100 μg Tx at P4 and dissected at P6. White arrows indicate AVMs. (E-G) Quantification of AVM number. n = 6-11 mice per group. Error bars: SEM. **** P-value < 0.0001, two-tailed unpaired t-test. (H-K) Vascular labeling with latex dye (red) of retinal and brain vessels in *Alk1^f/f^* (H and I) and *Alk1^f/f^ Mfsd2a Cre^ERT2^* (J and K) P6 pups. White arrows indicate AVMs. (L) Schematic representation of the experimental strategy used to delete *Alk1* in mice (P1-P6). Arrowheads indicate injection of 100 μg Tx at P1, P2 and P3 in *Alk1^f/f^ Esm1* and *Bmx Cre^ERT2^ mTmG* pups. (M and N) IB4 (blue) and GFP (white) staining of retinal flat mount from *Alk1^f/f^ Esm1 Cre^ERT2^ mTmG* (M) and *Alk1^f/f^ Bmx Cre^ERT2^ mTmG* (N). Scale bars: 500 μm (B-D, M-N), 200 μm (H and J), 2 mm (I and K).

Next, we examined the survival rate of these three mouse lines. *Alk1^f/f^ Mfsd2a Cre^ERT2^* mice died 5 to 6 days after gene deletion, most likely from ruptured brain AVMs, while *Alk1^f/f^ Bmx Cre^ERT2^* mice lived at least 50 days after gene deletion (Figure 3 A). Interestingly, the *Alk1^f/f^ Esm1 Cre^ERT2^* mutants died 10-11 days after Tx injection (Figure 3 A), suggesting they might develop AVMs in other tissues. Autopsy revealed massive intestinal hemorrhages in *Alk1^f/f^ Esm1 Cre^ERT2^* mice as a likely cause of death (Figure 3 B). To define the Esm1 expression in intestines, Tx was injected at P4 and *Esm1 Cre^ERT2^ mTmG* mice were analyzed at P14. GFP-positive cells were found in scattered capillaries of the mesenteries, the intestinal wall and the intestinal villi (Figure 3 C-D). We performed immunostaining of VE-Cadherin (VE-Cad) and GFP in P14 *Alk1^f/f^* and *Alk1^f/f^ Esm1 Cre^ERT2^ mTmG* mice (Figure 3 E-F). *Alk1^f/f^ Esm1 Cre^ERT2^ mTmG* developed GFP positive vascular malformations in capillaries of the intestinal villi (Figure 3 F). Injection of latex dye confirmed the presence of AVMs in the intestinal villi and in the mesenteries (Figure 3 G-J). To identify the presence of AVMs in other vascular beds, immunostaining and latex red dye injections were performed in P12 and P14 *Alk1^f/f^* and *Alk1^f/f^ Esm1 Cre^ERT2^ mTmG* mice (Figure 3 K-P). *Alk1^f/f^ Esm1 Cre^ERT2^ mTmG* developed GFP positive vascular malformations in retinal capillaries, and migration of Alk1 mutant tip cell progeny into the arteries was perturbed (Figure 3 K-L). Latex injection confirmed abnormal patterning of distal retinal arteries derived from the ESM1+ tip cells (Figure 3 M-N). The latex also revealed vascular malformations in the pial arteries of the brain in *Alk1^f/f^ Esm1 Cre^ERT2^* mice (Figure 3 O-P), but full-blown AVMs were not observed in retina or brain. These data indicate that loss of ALK1 signaling in ESM1 expressing capillaries leads to intestinal vascular malformations.

**Figure 3.**
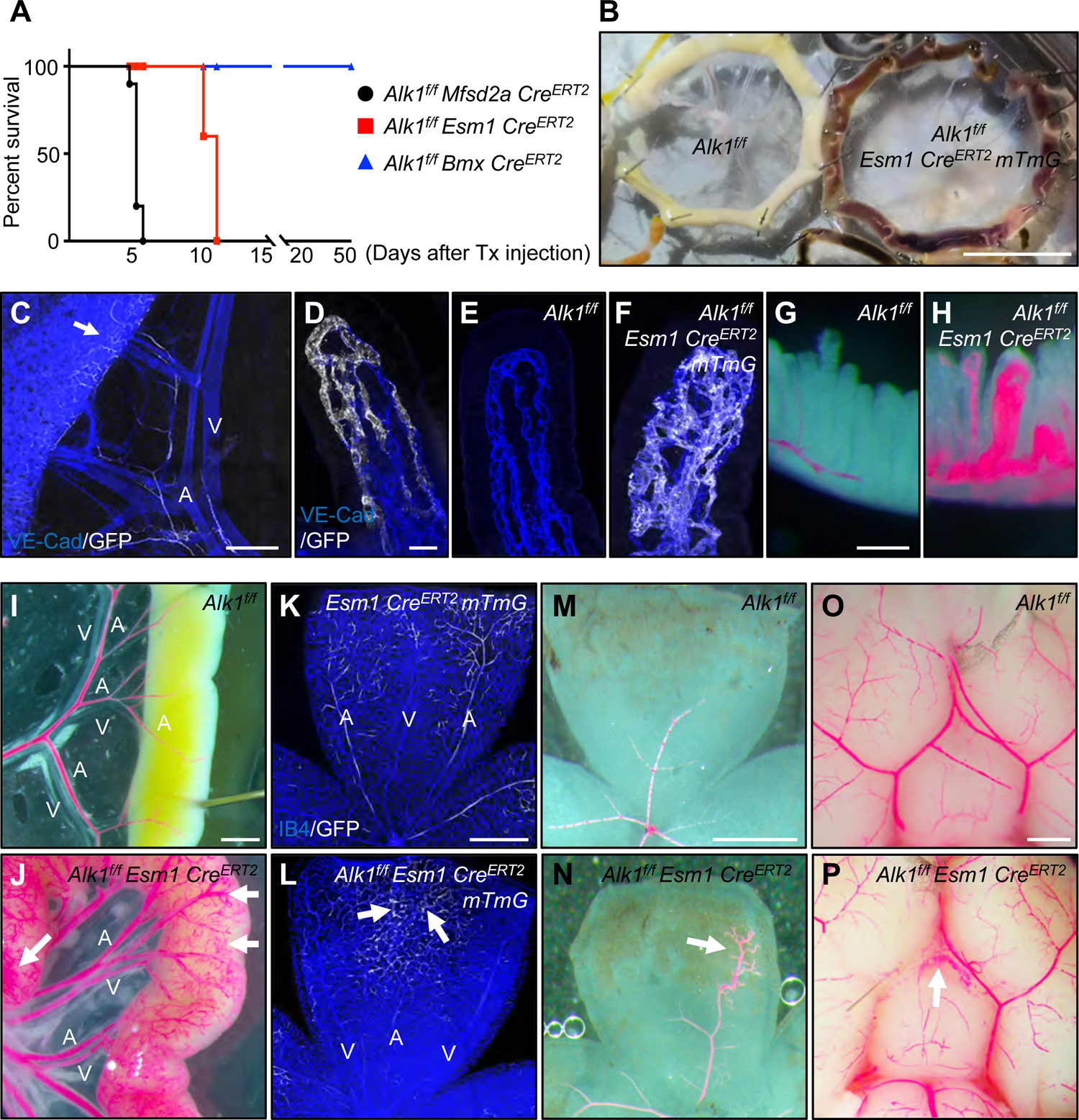
*Alk1^f/f^ Esm1 Cre^ERT2^ mTmG* mice display vascular malformations. (A) Survival curves for *Alk1^f/f^ Mfsd2aCre^ERT2^*, *Alk1^f/f^ Esm1Cre^ERT2^* and *Alk1^f/f^ BmxCre^ERT2^* mice injected with 100 μg Tx at P4. n = 8-10 mice/group. (B) Freshly dissected small intestines from P14 mice with the indicated genotypes after 100 μg Tx injection at P4. *Alk1^f/f^ Esm1 Cre^ERT2^ mTmG* mice displayed intestinal hemorrhage. (C and D) GFP (white) and VE-Cad (blue) staining of mesentery and gastrointestinal (GI) tract (C) and lacteals (D) from P14 *Esm1 Cre^ERT2^ mTmG*. 100 μg Tx was injected at P4. An arrow indicates *Esm1* positive capillary ECs (C). (E and F) VE-Cad (blue) and GFP (white) staining of jejunum lacteals from P14 *Alk1^f/f^* (E) and *Alk1^f/f^ Esm1 Cre^ERT2^ mTmG* (F). (G-J and M-P) 100 μg Tx was injected at P4 and dissected at P12. Vascular labeling with latex dye (red) of villi, GI tracts, retinas and brains in *Alk1^f/f^* (G, I, M and O) and *Alk1^f/f^ Esm1 Cre^ERT2^* (H, J, N and P) P12 pups. (K and L) 100 μg Tx was injected at P4 and dissected at P12 (K and L). IB4 (blue) and GFP (white) staining of retinal flat mounts from *Esm1 Cre^ERT2^ mTmG* (K) *and Alk1^f/f^ Esm1 Cre^ERT2^ mTmG* (L) P12 mice. An arrow indicates vascular malformations (J, L and N). A: artery, V: vein, Scale bars: 1 cm (B), 400 μm (C), 1 mm (I-J and M-P), 500 μm (K-N), 200 μm (G and H), 25 μm (D-F).

### Loss of ALK1 affects cell polarity and flow-migration coupling

To test if flow-mediated EC polarization was altered in the absence of ALK1, we dissected retinas at P6 48 h after Tx injection and immunolabeled with IB4 to detect ECs, DAPI to label nuclei, the Golgi marker GOLPH4 and ALK1 (Figure 4 A-F). To analyze the orientation of the Golgi toward the flow direction in the retinal vessels, we measured the angles between the EC nuclei and the Golgi as well as the predicted blood flow vectors (Figure 4 C’-F’, Figure 4 G). In *Alk1^f/f^* retinas, ALK1 expressing arterial, venous and capillary ECs polarized against the direction of blood flow (Figure 4 A, C, C’ and H). In contrast, ECs from *Alk1^f/f^ Mfsd2a Cre^ERT2^* retinas showed random Golgi distribution in veins, capillaries and AVMs (Figure 4 B, E, E’, F, F’ and H). Proximal arteries in *Alk1^f/f^ Mfsd2a Cre^ERT2^* retinas that maintained ALK1 expression were polarized normally against the flow (Figure 4B, D, D’ and H) Quantification of polarization using a polarity index (PI), which ranges from 1 (strongly polarized) to 0 (random distribution) confirmed that *Alk1^f/f^* retinal ECs were strongly polarized against the direction of blood flow, while *Alk1^f/f^ Mfsd2a Cre^ERT2^* mutant ECs in capillaries, veins and AVMs displayed poor polarization against the direction of blood flow (Figure 4 I). To determine whether these polarity defects preceded AVM development, Tx was injected at P4 and mice were analyzed after 24 h (P5) or 36 h (P5.5). Interestingly, AVMs started to appear at 24 h and were more pronounced at 36 h (Figure 4 J-K). Analysis of cell polarity in P5 *Alk1^f/f^ Mfsd2a Cre^ERT2^* mutants and controls showed that venous and capillary ECs from *Alk1^f/f^ Mfsd2a Cre^ERT2^* retinas displayed poorly polarized Golgi distribution (Figure 4 L, M), indicating that lack of flow-induced polarity preceded AVM formation and could be causally related to AVM development.

**Figure 4.**
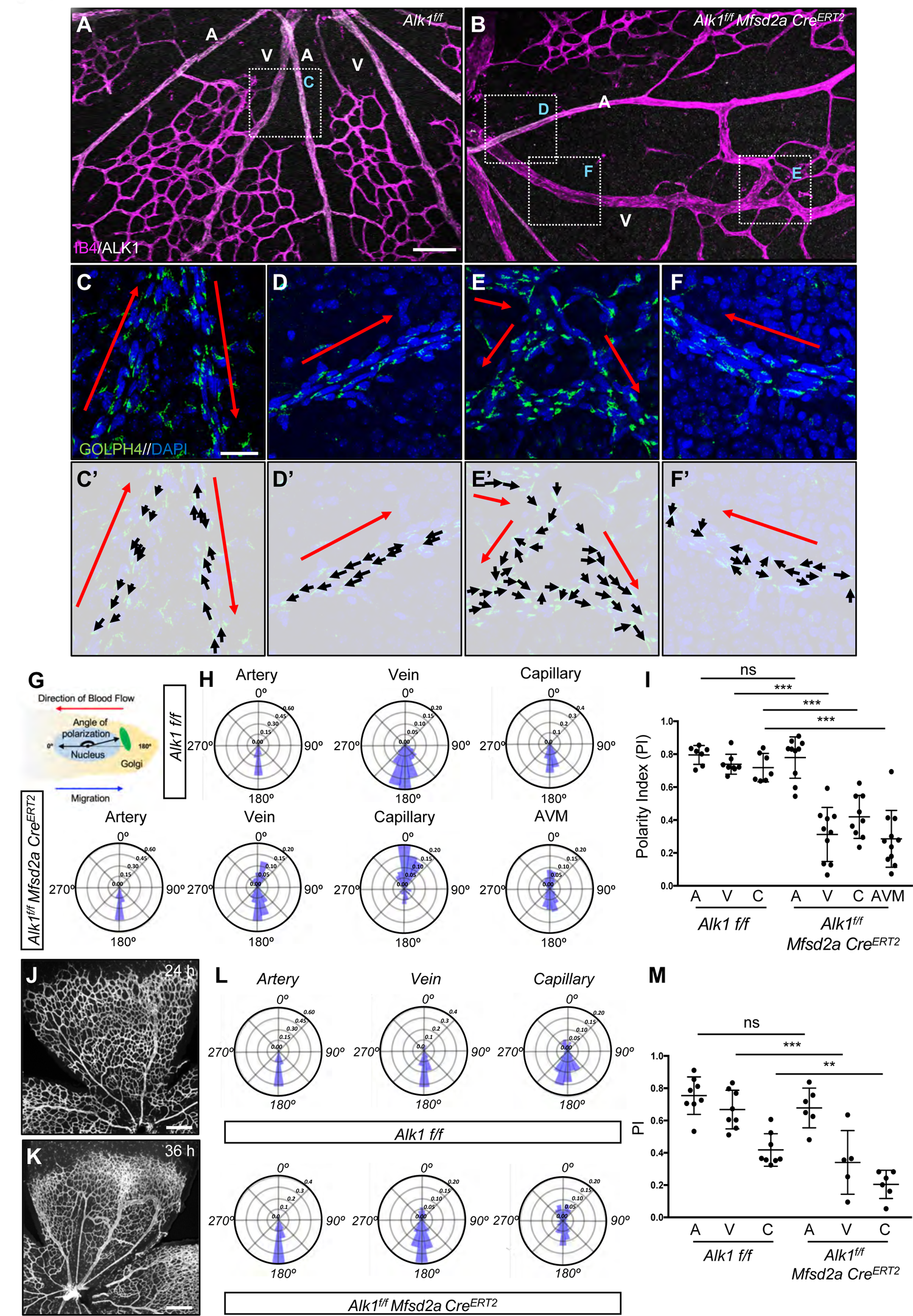
ALK1 controls cell polarization against the blood flow direction. (A-B) IB4 (Magenta) and ALK1 (white) staining of retinal flat mounts from *Alk1^f/f^* (A) and *Alk1^f/f^ Mfsd2a Cre^ERT2^* (B) pups injected with 100 μg Tx at P4 and dissected at P6. (C-F) Higher magnification of insets in A and B. GOLPH4 (green) and DAPI (blue) staining of retina flat mounts. Red arrows indicate the blood flow direction. (C’-F’) Background images from Figure 2 C-F and corresponding polarity vectors (black arrows). (G) The polarity axis of each cell was defined as the angle between the direction of blood flow and the cell polarity axis, defined by a vector drawn from the center of the cell nucleus to the center of the Golgi apparatus. (H) Angular histograms showing the distribution of polarization angles of ECs in the artery, vein and capillaries from *Alk1^f/f^* and artery, vein, capillary and AVM from *Alk1^f/f^ Mfsd2a Cre^ERT2^* mouse retinas. n = 7-11 retinas. (I) PI box plots of ECs from artery, vein and capillary from *Alk1^f/f^* and artery, vein, capillary and AVM from *Alk1^f/f^ Mfsd2a Cre^ERT2^* P6 retinas. n = 7-11 retinas. (J and K) IB4 (gray) staining of retinal flat mounts from *Alk1^f/f^ Mfsd2a Cre^ERT2^* pups injected with 100 μg at P4 and dissected after 24 h (P5) (J) and 36 h (P5.5) (K). (L) Angular histograms showing the distribution of polarization angles of ECs in the artery, vein and capillary from *Alk1^f/f^* and *Alk1^f/f^ Mfsd2a Cre^ERT2^* P5 retinas at 24 h after Tx injection. (M) PI box plots of ECs from artery, vein and capillary from *Alk1^f/f^* and *Alk1^f/f^ Mfsd2a Cre^ERT2^* retinas at 24 h after Tx injection. n = 5-8 retinas/group. Error bars: SEM. **P-value < 0.01, ***P-value < 0.001, ns: nonsignificant, two-tailed unpaired t-test. Scale bars: 100 μm (A-B), 20 μm (C-F) and 500 μm (J-K)

To explore whether laminar flow affected the polarization of *Alk1* mutant cells *in vitro*, we performed scratch wound assays with human umbilical vein endothelial cells (HUVECs) that were cultured in static conditions or subjected to laminar shear stress (15 dynes/cm^2^) and stained with a Golgi marker to determine cell polarity angles^22^. In static conditions, control siRNA transfected HUVECs were polarized towards the scratch areas in both the left and the right side of the wound (Fig.5A,C). Under laminar shear, the cells on the left side upstream of the scratch repolarize in the opposite direction to align against the flow (Fig.5E,G). By contrast, *ALK1* deficient HUVECs showed random polarization in static conditions (Fig.5B,D). Most strikingly, they were unable to polarize against the direction of flow in the upstream scratch areas, and even the downstream polarization against the flow was impaired (Figure 5 F,H). Polarity index calculation showed that flow significantly enhanced polarization of control siRNA transfected cells, and that *ALK1* deletion prevented flow induced polarization (Figure 5 I).

**Figure 5.**
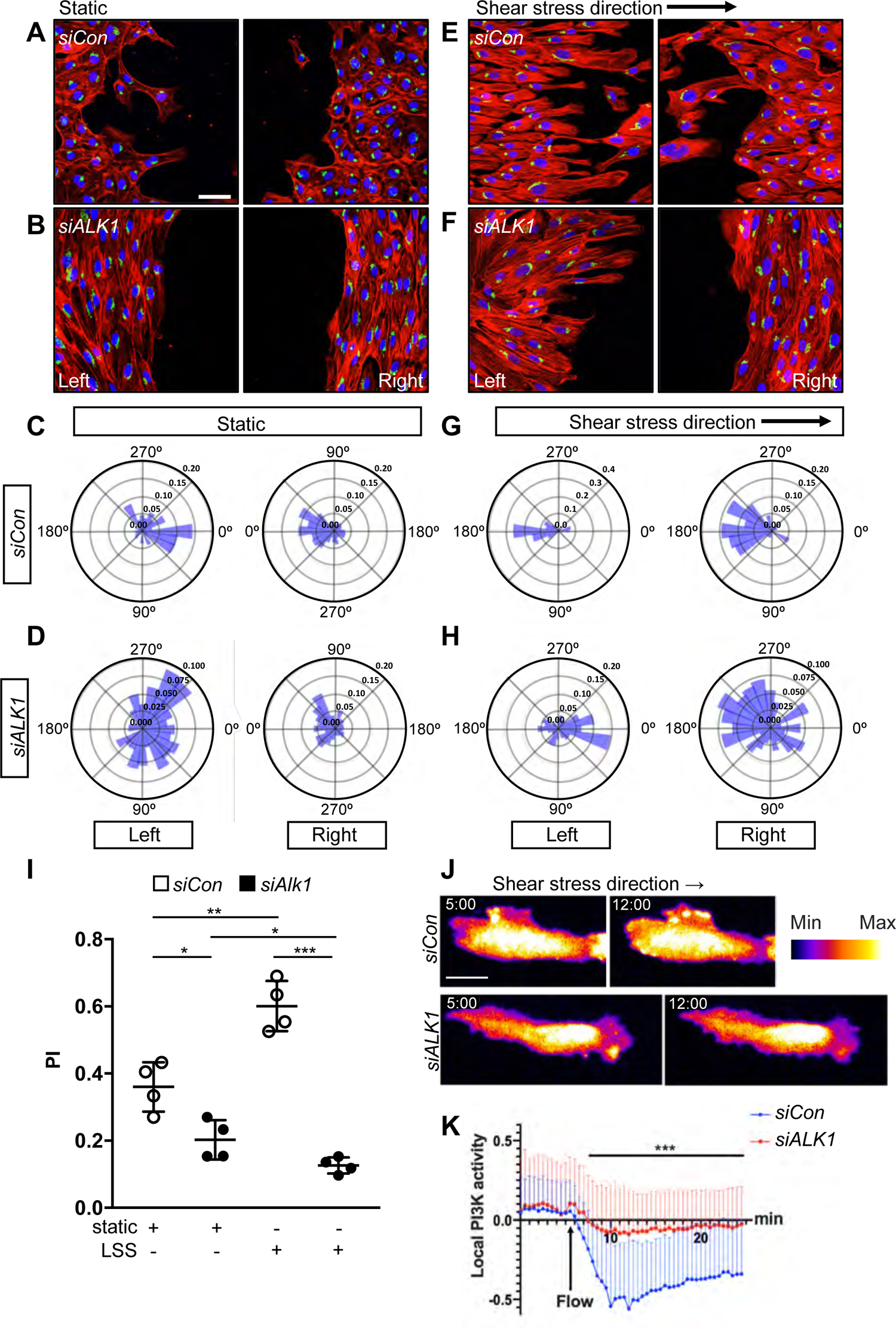
ALK1 controls EC polarization against the flow direction *in vitro*. (A-B) Representative images of wound-healing assays after 18 h showing polarity angles of HUVECs transfected with Control (*siCon*) (A) or *ALK1* (*siALK1*) (B) siRNAs under static conditions and immunolabeled with phalloidin(red), GM130 (green), and DAPI (blue). (E-F) Representative images of wound-healing assays showing polarity angles of *siCon* (E) or *siALK1* (F) HUVECs with 18 h exposure to laminar shear stress (LSS) at 15 dynes/cm^2^. Left panels are upstream and right panels are downstream of flow. (C-D and G-H) Angular histograms showing polarization angles of *siCon* (C and G) or *siALK1* ECs (D and H) at 18 h after scratch with (G-H) or without (C-D) LSS. Left is upstream and right is downstream of flow (G and H). (I) PI box plots of upstream (left) scratch areas from *siCon* or *siALK1* transfected HUVECs at 18 h after with or without LSS. (C-I) n=6-8 images from 3 independent experiments. Error bars: SEM. *P-value < 0.05, **P-value < 0.01, ***P-value < 0.001, two-tailed unpaired t-test. (J) Representative time lapse images of *siCon* or *siALK1* HUVECs stably transduced with PH-AKT-mClover3 and plasma membrane targeting sequence of LCK-mRuby3. HUVEC monolayers in microfluidic chambers were exposed to 12 dynes/cm^2^ LSS under the microscope. 5 min (static) and 12 min (LSS) images were selected from the movies. The surface is color-coded by the value of PH-AKT intensity. (K) Local activation of PI3K was quantified by image analysis. PH-AKT intensity was normalized with average static intensity at each time point. 0 - 5 min: static and 5 - 24.5 min: LSS, n= 61, 41 cells from 3 independent experiments, Error bar: SEM. ***P-value < 0.001, two-tailed unpaired t-test. Scale bars: 50 μm (A-B and E-F), 20 μm (J).

Flow induces localization of phosphorylated AKT to the upstream edge of ECs^30^. Pleckstrin homology domain of AKT fused to GFP (PH-AKT-GFP) is a well-established biosensor of PI3K local activity which shows plasma membrane localized PH-AKT-GFP upon shear stress^31^. To examine whether ALK1 affected flow-induced PI3K localization, we performed live cell imaging. PH-AKT-mClover3 together with plasma membrane marker (LCK-mRuby3) were co-expressed as a biosensor and an internal control respectively. Control siRNA transfected HUVECs showed polarized activation of PI3K on their upstream edge within 5 minutes after flow, consistent with upstream Golgi polarization. By contrast, *ALK1* knockdown significantly diminished this effect (Figure 5 J and K, Supp. Figure 3 movies). This result indicated that ALK1 is required for flow-induced PI3K-AKT polarization.

### Blockade of integrin prevents AVM formation

Golgi orientation into scratch wounds and under flow is driven by integrin binding to ECM proteins and signaling to CDC42^32, 33^. Additionally, integrin activation and signaling is modulated by VEGFR2-PI3K signaling, which is altered following ALK1 deletion^12, 13, 18^ ^34^. VEGFR2 interacts with integrins αvβ3 and α5β1 during vascularization^35–37^, prompting us to test if VEGFR2-integrin signaling was enhanced in *ALK1* deficient ECs. Interestingly, immunolabeling with antibodies recognizing integrin β1 (ITGB1), α5 (ITGA5) and αv (ITGAV) showed increased ITGB1, ITGA5 and ITGAV expression in the AVM areas of P8 *Alk1^f/f^ Mfsd2a Cre^ERT2^* retinas when compared to wildtype controls (Figure 6 A-C and Supp. Figure 4 A-C). We next tested if the interaction of VEGFR2 and integrin was affected in *ALK1* deleted cells. While total levels of ITGB1, ITGA5 and ITGAV were moderately increased in *ALK1* knockdown cells, their co-immunoprecipitation with VEGFR2 was enhanced to a greater degree (Figure 6 D,E). These results suggested that targeting integrins signaling with inhibitors could rescue AVM formation. To test this idea, we administered Cilengitide, a small molecule inhibitor for integrin αvβ3 and αvβ5, or ATN161, a peptide inhibitor for integrin α5β1. *Alk1* deletion was induced by Tx injection at P4, inhibitors were given i.p at 5 mg/kg at P4 and P5, and mice were analyzed at P6 (Figure 6 F). Both Cilengitide and ATN161 decreased AVM formation in *Alk1^f/f^ Mfsd2a Cre^ERT2^* or *Alk1^f/f^ Cdh5 Cre^ERT2^* mice (Figure 6 G-N). Immunostaining of Golgi markers showed that integrin inhibitors rescued polarization of *Alk1* mutant cells against the direction of blood flow (Figure 6 O-S).

**Figure 6.**
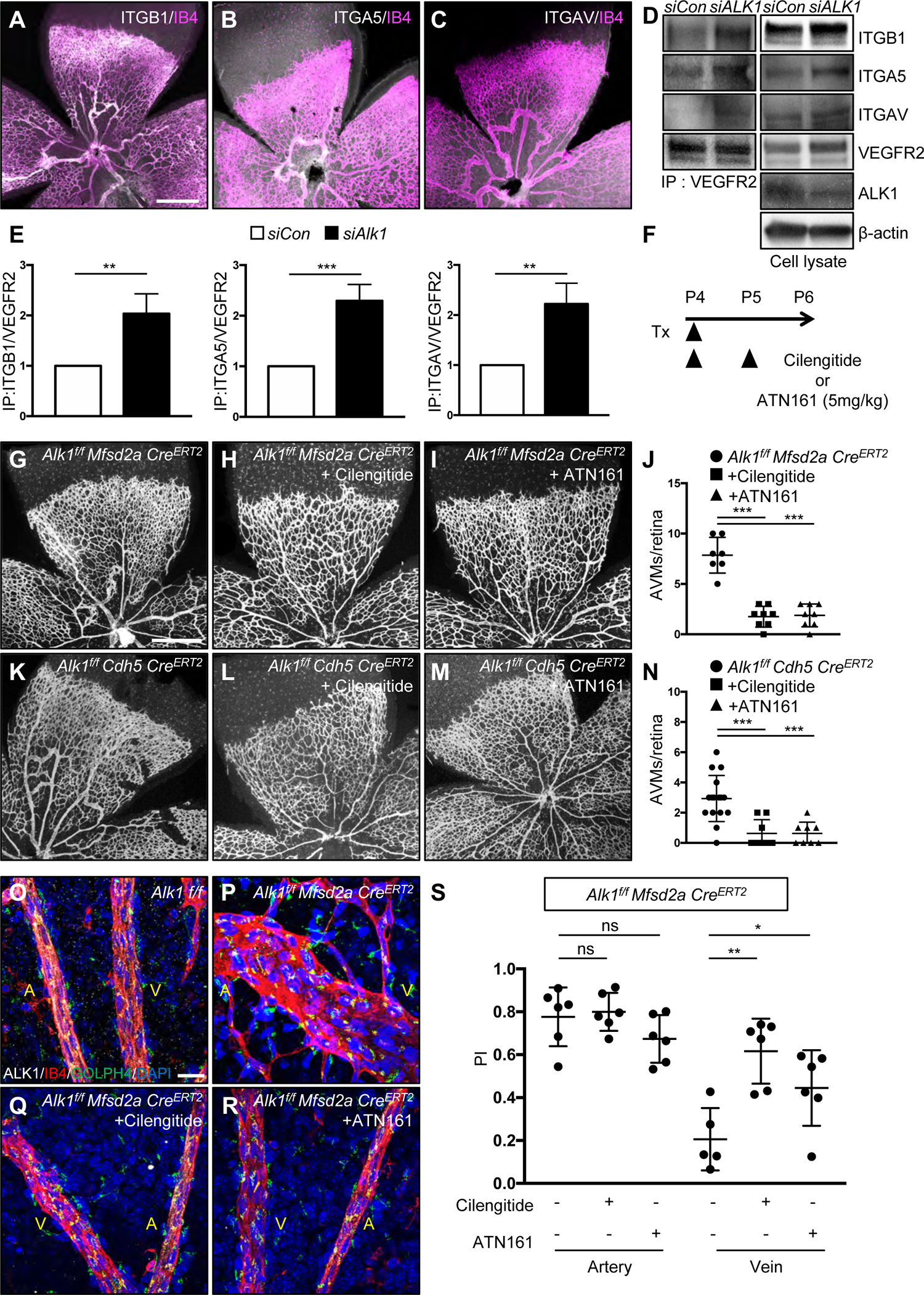
Integrin inhibition prevents AVM formation in *Alk1* mutant retinas. (A-C) IB4 (Magenta) and ITGB1(A, white), ITGA5 (B, white) or ITGAV (C, white) staining of retinal flat mounts from P8 *Alk1^f/f^ Mfsd2a Cre^ERT2^* pups. (D) VEGFR2 immunoprecipitation in *siCon* or *siALK1* HUVECs and western blot analysis for ITGB1, ITGA5 and ITGAV. VEGFR2, ITGB1, ITGA5, ITGAV, ALK1 and β-actin expression from the total cell lysates are shown as loading controls. (E) Quantification of ITGB1, ITGA5 or ITGAV levels normalized to VEGFR2 from immunoprecipitation. **P<0.01, ***P-value < 0.001, two-tailed unpaired t-test. (F) Experimental strategy to assess the effects of integrin inhibitors in *Alk1* deleted retinas. Arrowheads indicate the time course of Tx (100 μg) and Cilengitide (5mg/kg), ATN161 (5mg/kg) or vehicle administration. (G-I and K-M) IB4 staining of P6 retinal flat mounts from *Alk1^f/f^ Mfsd2a Cre^ERT2^* (G-I) or *Alk1^f/f^ CDH5 Cre^ERT2^* (K-M) injected with Cilengitide (H and L) or ATN161 (I and M) at P4 and P5. (J and N) Quantification of the AVM number. Each dot represents one retina. n = 7-16 retinas per group. Error bars: SEM. ***P-value < 0.001, One-way ANOVA with Holm-Sidak test. (O-R) IB4 (Magenta), Alk1 (white), GOLPH4 (green) and DAPI (blue) staining of retina flat mounts from *Alk1^f/f^* (O), *Alk1^f/f^ Mfsd2a Cre^ERT2^* (P), Cilengitide (Q) or ATN161 (R) injected *Alk1^f/f^ Mfsd2a Cre^ERT2^* pups. A: artery, V: vein, (S) PI box plots of ECs from artery and vein from *Alk1^f/f^*, *Alk1^f/f^ Mfsd2a Cre^ERT2^*, Cilengitide or ATN161 injected *Alk1^f/f^ Mfsd2a Cre^ERT2^* retinas. n=5-8 retinas/group. Error bars: SEM. *P-value < 0.05, **P-value < 0.01, ns: nonsignificant, One-way ANOVA with Holm-Sidak test. Scale bars: 500 μm (A-C, G-I and K-M), 20 μm (O-R).

### ALK1 controls integrin mediated Hippo pathway signaling

Previous data reported an interaction between BMP9/ALK1 signaling and the YAP/TAZ pathway, and both of these pathways are regulated by blood flow *in vivo*^38, 39^. Moreover, integrins are potent regulators of YAP/TAZ activation in many systems including ECs^40–42^. Activation of YAP/TAZ results in both protein stabilization and nuclear translocation, with induction of target gene expression. To test whether laminar shear and ALK1 affected integrin and YAP/TAZ protein expression, HUVECs were transfected with control or *ALK1* siRNA and cultured in static conditions or under laminar shear stress (15 dynes/cm^2^) for 18 h. Protein extracts from these cells were analyzed by Western blot with antibodies against integrins, YAP or TAZ and expression levels were compared to b-actin. Interestingly, ITGB1, ITGA5 and ITGAV as well as YAP and TAZ were all significantly increased in *ALK1* deleted ECs when compared to control siRNA transfected cells, and their expression was further increased in laminar shear stress conditions (Figure 7 A-B). YAP and TAZ protein expression was also greatly increased and appeared more nuclear in *Alk1^f/f^ Mfsd2a Cre^ERT2^* retina AVMs when compared to *Alk1^f/f^* control ECs (Figure 7 C-F). Immunostaining of *ALK1* deficient HUVECs with YAP and TAZ antibodies confirmed an increase of YAP/TAZ expression in *ALK1* siRNA transfected HUVECs, and moreover revealed that YAP and TAZ located both in the cytosol and the nucleus *ALK1* deficient cells, while they were located in the cytosol of control siRNA treated ECs (Figure 7 G-H). To test whether other HHT pathway components *ENG* and *SMAD4* also affected YAP/TAZ activity, we deleted *ALK1*, *SMAD4*, and *ENG* in HUVECs and immunostained for YAP and TAZ (Figure 8 A). We found that YAP and TAZ showed translocation to the nucleus in *ALK1*, *SMAD4*, and *ENG* deleted ECs (Figure 8 A). Furthermore, we found that the YAP/TAZ inhibitor Verteporfin (VP) blocked YAP/TAZ activity and translocation to the nucleus in *ALK1*, *SMAD4*, and *ENG* depleted HUVECs (Figure 8 A). To examine whether the inhibition of YAP/TAZ could improve AVMs *in vivo*, we first administered VP (50 mg/kg, i.p.) into P4 and P5 *Alk1^f/f^* control mice (Figure 8 B). VP injected control retina developed blunted endothelial tip cells at the angiogenic front (Figure 8 C), as reported in genetically YAP/TAZ deficient endothelial mouse retinas^38, 43, 44^ indicating that the pharmacological inhibition was effective. Next we injected VP into *Alk1^f/f^ Cdh5 Cre^ERT2^* or *Alk1^f/f^ Mfsd2a Cre^ERT2^* retinas, which led to a significant reduction of AVM formation and hemorrhage when compared to DMSO vehicle treated mutant retinas (Figure 8 C-E). VP treatment also rescued the polarization against the direction of blood flow (Figure 8 F-G). These results demonstrated that ALK1 regulates Hippo pathway activation, and that VP-mediated inhibition of YAP/TAZ nuclear translocation improved flow migration coupling and prevented AVMs in *Alk1* mutant ECs.

**Figure 7.**
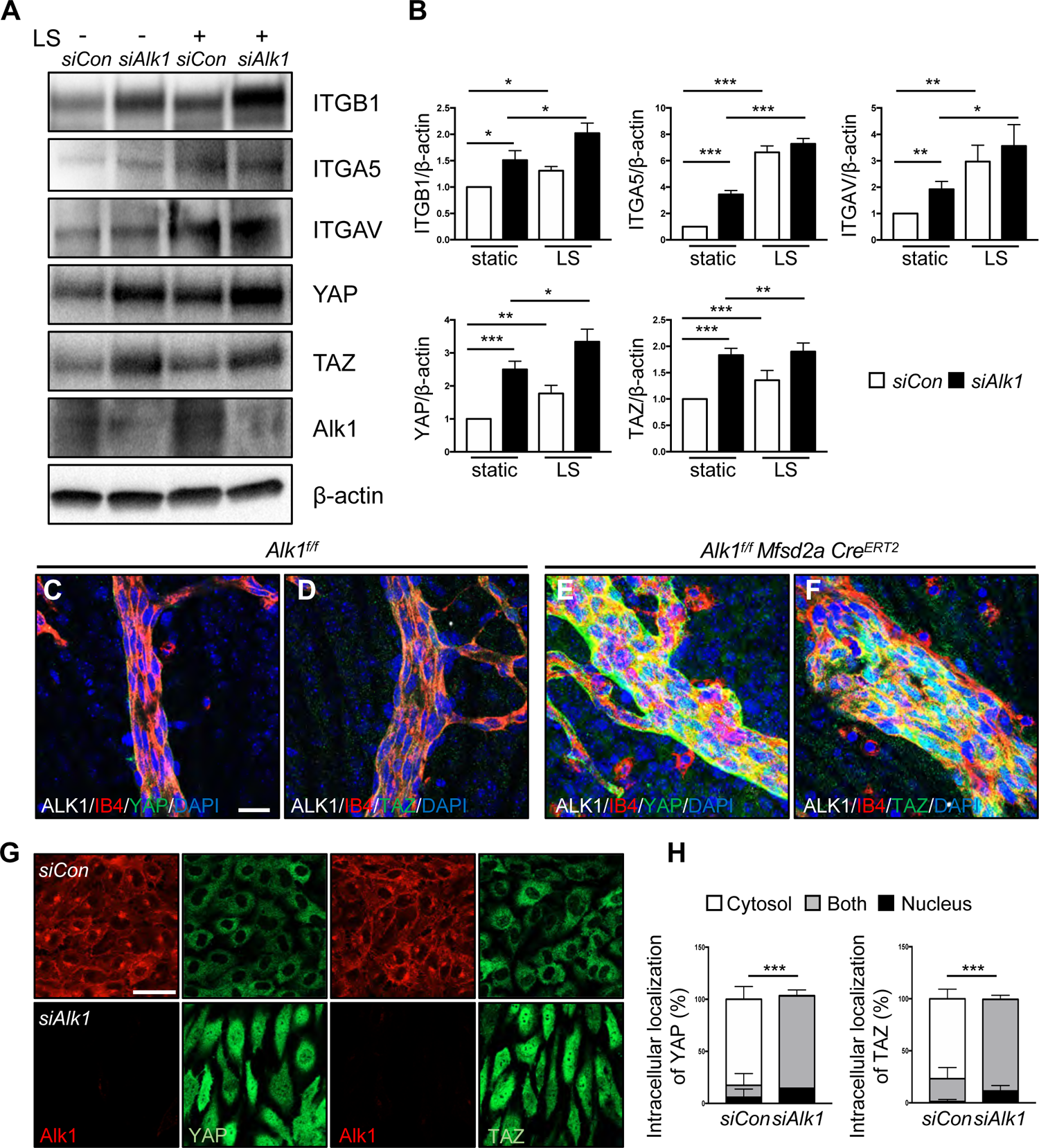
ALK1 controls YAP/TAZ expression and localization. (A) Western blot analysis of HUVECs transfected with control and *ALK1* siRNAs followed by 18 h exposure to LSS (15 dynes/cm^2^). (B) Quantification of ITGB1, ITGA5, ITGAV, YAP or TAZ levels normalized to β-actin. *P<0.05, **P<0.01, ***P<0.001, two-tailed unpaired t-test. (C-F) YAP and TAZ (green), ALK1 (gray), IB4 (red), DAPI (blue) staining of retinal flat mounts from P8 Alk1^f/f^ (C-D) or Alk1^f/f^ Mfsd2aCre^ERT2^ (E-F) pups. A scale bar: 20 μm (A-F) (G) YAP or TAZ (green) and Alk1 (red) staining of *siCon* and *siALK1* HUVECs. A scale bar: 50 μm. (H) Quantification of YAP and TAZ localization from *siCon* and *siALK1* transfected HUVECs. ***P<0.001, n = 3 independent experiments. Multiple comparisons with Holm-Sidak test.

**Figure 8.**
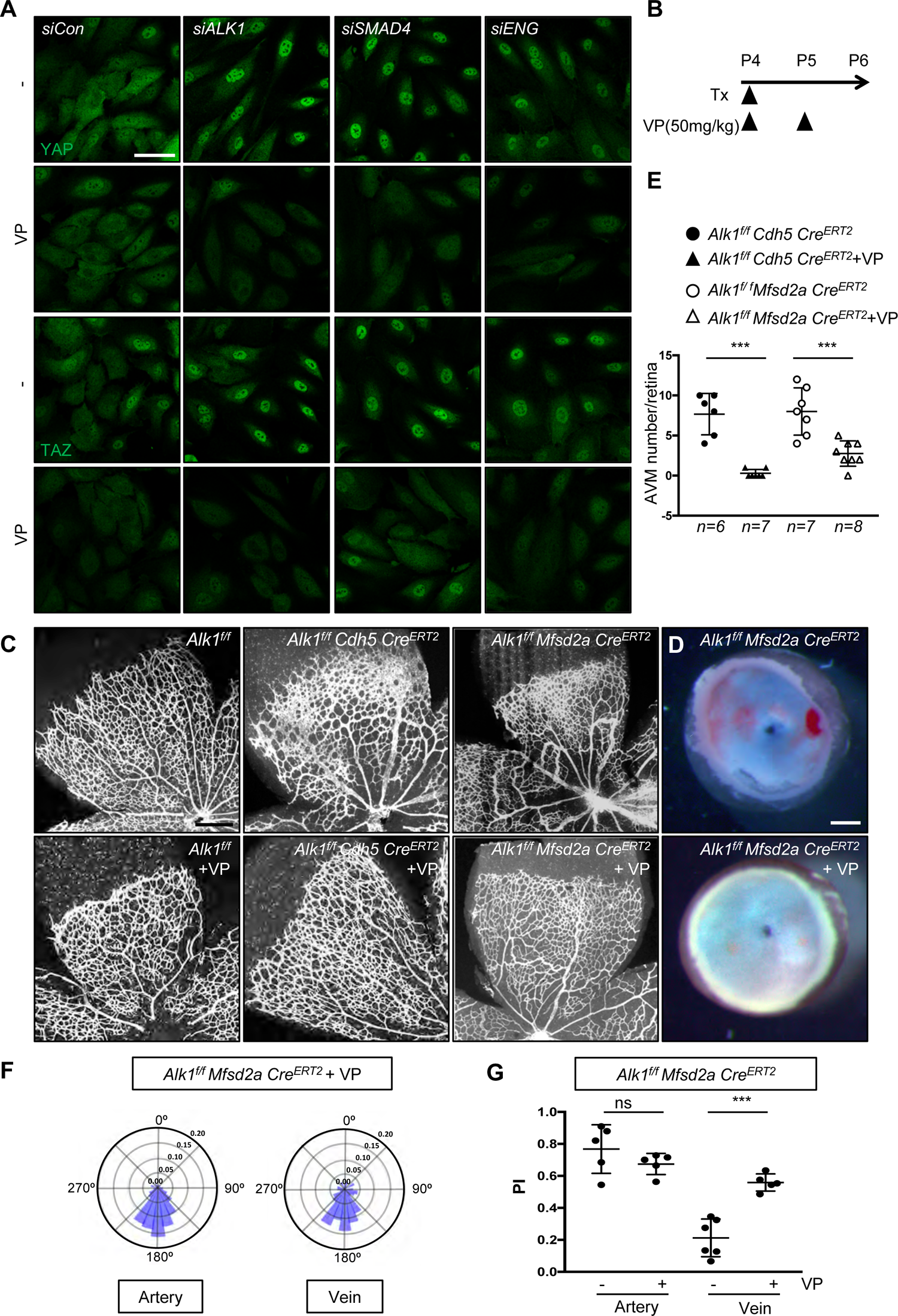
YAP/TAZ inhibition improves AVM formation in *Alk1* mutant retinas. (A) YAP and TAZ staining of *siCon*, *ALK1*, *SMAD4* or *ENG* siRNAs transfected HUVECs treated with DMSO or Verteporfin (VP, 5 μM) for 6 h. Nuclear YAP/TAZ localization in *siALK1*, *siSMAD4* or *siENG* ECs is blocked by VP treatment. A scale bar: 50 μm. (B) Experimental strategy to assess the effects of YAP/TAZ inhibition in EC specific *Alk1* deleted vasculature. Arrowheads indicate the time course of Tx (100 μg) and VP (50mg/kg) or vehicle administration. (C) IB4 staining of P6 retinal flat mounts from VP injected *Alk1^f/f^*, *Alk1^f/f^ CDH5 Cre^ERT2^* or *Alk1^f/f^ Mfsd2a Cre^ERT2^* mice. (D) Stereomicroscopy images of vehicle or VP injected *Alk1^f/f^ Mfsd2a Cre^ERT2^* retinas. (E) Quantification of the AVM number/retina. Each dot represents one retina. n = 6-8 retinas per group. Error bars: SEM. ***P-value < 0.001, two-tailed unpaired t-test. (F) Angular histograms showing polarization angles of artery and vein from *Alk1^f/f^ Mfsd2a Cre^ERT2^* with VP. (G) PI box plots of *Alk1^f/f^ Mfsd2a Cre^ERT2^* with vehicle or VP. n=5-6 retinas, Error bars: SEM, ***P-value < 0.001, ns: nonsignificant, two-tailed unpaired t-test. Scale bars: 50 μm (A), 500 μm (C), 300 μm (D)

### integrin and PI3K function upstream of YAP/TAZ

To elucidate whether ALK1 modulation of the Hippo pathway was integrin and PI3K dependent, we examined YAP/TAZ activity and translocation to the nucleus upon integrin inhibitor or PI3K inhibitor treatment. Intriguingly, YAP/TAZ activity and nuclear translocation was significantly reduced in *ALK1* deleted HUVECs treated with integrin inhibitors Cilengitide or ATN161 (Figure 9 A and B), and in *ALK1* deleted cells treated with the PI3K inhibitor wortmannin (Supp. Figure 5 A). Next, we injected Tx into P4 *Alk1^f/f^ Mfsd2a Cre^ERT2^* mice and administered Cilengitide and ATN161 at P4 and P5 to examine the YAP/TAZ activity and nuclear translocation. Interestingly, YAP/TAZ nuclear translocation was blocked in integrin inhibitor injected *Alk1^f/f^ Mfsd2a Cre^ERT2^* retinas when compared to *Alk1^f/f^* control retinas (Figure 9 C).

**Figure 9.**
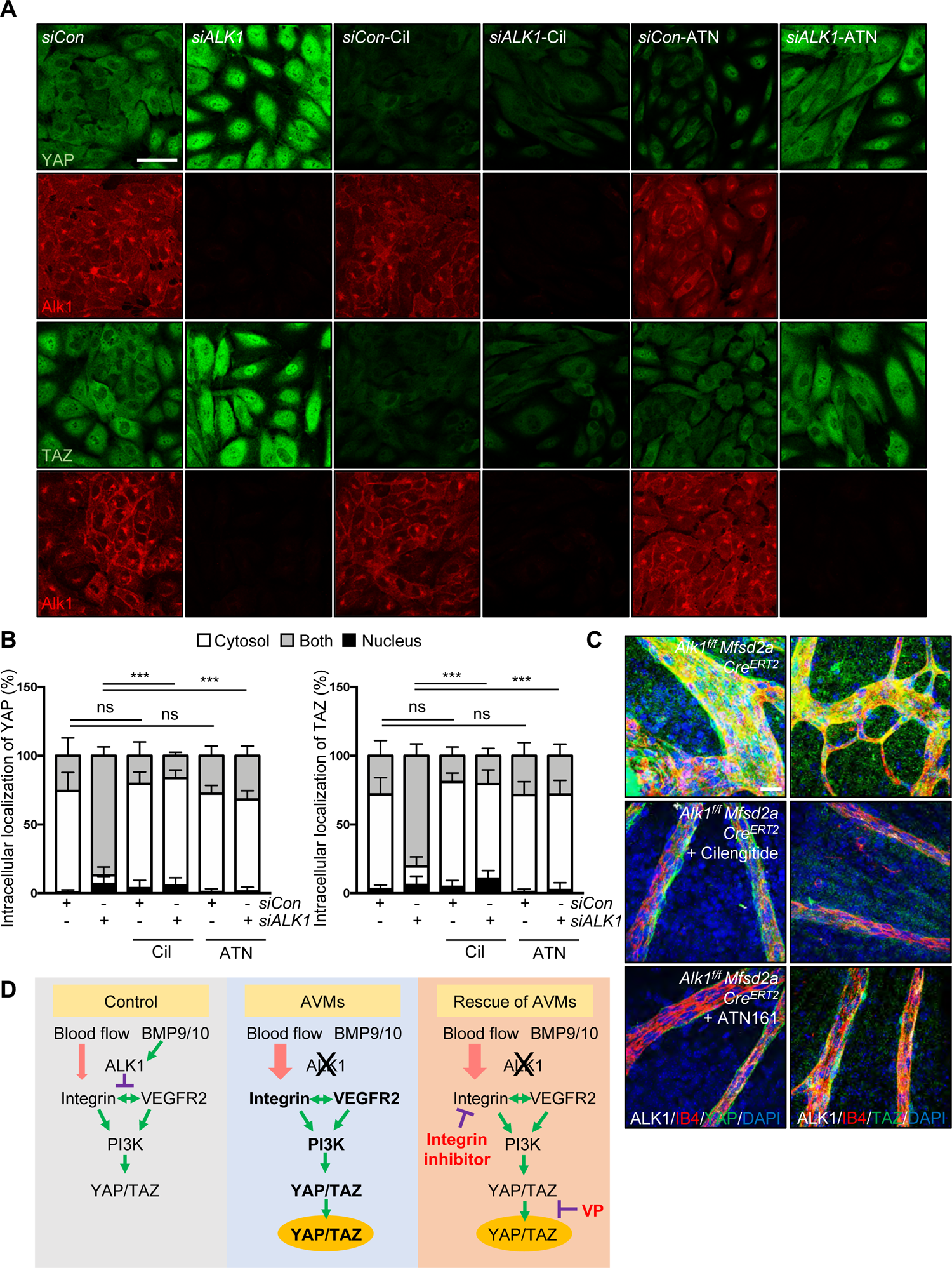
Integrin acts upstream of YAP/TAZ in an ALK1 dependent manner. (A) YAP and TAZ staining for control, *ALK1* siRNAs transfected HUVECs treated with PBS, Cilengitide (Cil, 5 μM) or ATN161 (ATN, 5 μM) for 12 h. Nuclear YAP/TAZ localization in *siALK1* ECs is blocked by Cilengitide and ATN161 treatment. (B) Quantification of YAP and TAZ localization from *siCon* and *siALK1* transfected HUVECs. ***P<0.001, n = 3 independent experiments. Error bars: SEM. ***P-value < 0.001, ns: nonsignificant, Multiple comparisons with Holm-Sidak test. (C) YAP and TAZ (green), ALK1 (white), IB4 (red) and DAPI (blue) staining of retinal flat mounts from Cilengitide or ATN161 injected *Alk1^f/f^ Mfsd2a Cre^ERT2^* P6 mice. (D) A model for ALK1-integrin-YAP/TAZ signaling in maintenance of vascular quiescence. In quiescence, ALK1 signaling represses PI3K activation downstream of integrin-VEGFR2 signaling, through inhibition of YAP/TAZ expression and localization. ALK1 deletion results in increased integrin-VEGFR2 signaling, and consequently in excessive YAP/TAZ expression and localization to the nucleus, thereby inducing vascular defects. Blocking integrin-ECM interaction with integrin inhibitors or YAP/TAZ localization with YAP/TAZ inhibitor rescues vascular malformations in *Alk1* deficient mice. Scale bars: 50 μm (A), 20 μm (C)

## Discussion

This study extends our previous knowledge by identifying the origin of AVM forming cells from capillaries and veins and the role of integrin and Hippo pathway signaling in HHT. The data are consistent with a model whereby the presence of ALK1 suppressed integrin-VEGFR2 signaling interactions, which limited downstream PI3K activation and signaling to YAP/TAZ, as indicated by changes in protein levels and nuclear translocation. In the absence of ALK1, enhanced VEGFR2-integrin-PI3K signaling stabilized YAP/TAZ and promoted nuclear translocation. Pharmacological inhibition of integrin or YAP/TAZ signaling prevented vascular malformations in *Alk1* deficient mice (Figure 9 D).

In retinal development, *Mfsd2a*-positive capillary-venous as well as *Esm1*-positive tip cells migrated against the blood flow direction towards retinal arteries. This is consistent with work published by others and highlights endothelial flow-migration coupling as a critical process driving vascular remodeling^22, 45, 46^. The current concept suggests that, in response to blood flow, ECs migrate from low flow segments (veins and capillaries) towards high flow segments (arteries)^45^. In this study, we tested the hypothesis that disruption of flow-migration coupling and resulting accumulation of ECs in capillaries could cause capillary enlargement and thereby precipitate AVM formation. Consistent with this model, deletion of *Alk1* in capillaries and veins using *Mfsd2a Cre^ERT2^* led to disruption of Golgi polarization against the flow direction and caused retinal and cerebral AVMs, while arterial-specific *Alk1^f/f^ Bmx Cre^ERT2^* mice developed no AVMs. Mfsd2a is a brain-specific endothelial gene^25, 47^, hence our analysis of these mice was restricted to the brain and retina. In addition, a recent study showed that capillary/venous-specific deletion of the ALK1 co-receptor ENG using *ENG^fl/fl^ Apj-Cre^ERT2^* mice induced retinal AVMs^48^, indicating that the Alk1-ENG complex is required in capillaries and veins to prevent AVM formation, and that defective flow-migration coupling is a hallmark of HHT. Whether venous or capillary ECs, or both, are involved in the vascular malformations, needs to be further investigated and will require generation of capillary or vein-specific *Cre* driver lines.

Interestingly, deletion of *Alk1* in *Esm1*-positive tip cells also led to defective flow-migration coupling and accumulation of the mutant cells in the vascular plexus ahead of the arteries, while control cells colonized the arterial tree. This underscores an important role of *Alk1* in flow-migration coupling of retinal tip cells, but produced only mild retinal and brain vascular malformations when compared to pan-endothelial *Alk1^f/f^ Cdh5 Cre^ERT2^* and *Alk1^f/f^ Mfsd2a Cre^ERT2^* mice. One possible reason for the discrepant phenotypes are different flow environments: *Esm1*-positive tip cells migrate in a low-flow environment, whereas AVMs develop in high-flow regions of the retina, close to the optic nerve, and we and others have previously reported that blood flow potentiates ALK1-ENG-mediated shear stress sensing^15, 27^.

Quite strikingly, despite the lack of AVMs in retina and brain, the *Alk1^f/f^ Esm1 Cre^ERT2^* mice developed intestinal AVMs and succumbed to intestinal hemorrhage. Analysis of *Esm1*-driven GFP labeling revealed expression in capillary endothelium of the mesenteries, the gut wall and the intestinal villi, and GFP positive cells formed AVMs in those regions in *Alk1^f/f^ Esm1 Cre^ERT2^* mutant mice. Hence, capillary function of *Alk1* was required to prevent intestinal AVM formation. Further analysis is required to assess whether flow-migration coupling also underlies intestinal vascular remodeling, but such studies will require endothelial specific fluorescent Golgi reporter mice to determine endothelial cell polarity.

Our *in vitro* data revealed that loss of ALK1 displayed disrupted endothelial Golgi orientation and polarization against the blood flow direction. Blocking blood flow in zebrafish *alk1* mutants prevented AVM formation, directly demonstrating that blood flow induces AVM formation in the absence of ALK1^49^. We and others have previously reported that BMP9/10-Alk1 signaling mechanistically links flow sensing and VEGFR2-PI3K/AKT pathway activation ^12–15, 17, 20, 50^. Alk1 signaling counteracted both flow and growth factor-induced AKT activation, and the absence of Alk1 overactivated PI3K/AKT signaling in AVMs ^51–53^. We extend these findings here by demonstrating that ALK1 is required for flow-induced PI3K-AKT polarization against the direction of blood flow. Besides enhanced VEGFR2/PI3K signaling, another study showed that loss of SMAD4 increased Angiopoietin2 and decreased TIE2 receptor expression^54^. Blocking Angiopoietin2 prevented AVM formation and normalized vessel diameters in endothelial *Smad4* deficient mice^54^. But TIE2 accumulated within the AVMs^54^, suggesting that increased TIE2 signaling could contribute to enhanced PI3K signaling in AVMs.

As new mechanistic findings, we report that integrins and YAP/TAZ signaling are involved in ALK1 signaling and AVM formation. Integrins are heterodimeric transmembrane receptors that are activated by flow and then bind to specific ECM proteins. RGD peptides or neutralizing antibodies against integrin α_5_β_1_ prevented laminar shear stress-induced increase in EC adhesion^55–57^. This correlates with our data that *Alk1* mutant showed increased integrins in AVM regions and Cilengitide and ATN161 improved AVMs in *Alk1* mutant mice. Cilengitide is a selective α_v_β_3_ and α_v_β_5_ integrin inhibitor. Phase 3 trials using Cilengitide for glioblastoma patients have failed to improve patient survival, however they were well tolerated^58^. We reasoned that integrin antagonists could be a safe treatment to improve the AVMs in *Alk1* mutants, thereby identifying Integrin antagonists as viable candidates for therapy in HHT patients.

Hippo-YAP/TAZ signaling regulates organ size, tissue regeneration and self-renewal as well as vascular development^38, 39, 59–62^. Endothelial YAP/TAZ are important for the activation of CDC42 and for junction integrity and stabilization to control cell polarity^63^. In zebrafish, YAP translocates to the nucleus in response to blood flow and promotes cell migration and proliferation^39^, suggesting that nuclear YAP/TAZ accumulation in ALK1 deficient ECs could contribute to enhanced proliferation. One of the endothelial YAP/TAZ targets CCN1 increases activation of the VEGFR2 and PI3K signaling pathways by binding with integrin α_v_β_3_ and VEGFR2, creating a positive feedback loop that maintains endothelial polarity^64^. Moreover, YAP/TAZ can integrate mechanical signals with BMP signaling to maintain junctional integrity^38^. Alk1 appears to mainly affect nuclear Yap/TAZ function. We found that loss of ALK1 increased protein levels and co-immunoprecipitation of integrins and VEGFR2 in a flow-dependent manner, leading to overactivated and mislocalized PI3K signaling and increased YAP/TAZ nuclear localization. Nuclear Yap/TAZ could complex with TEAD and transcriptionally regulate YAP/TAZ target gene expression including integrin ligands such as CYR61 and CTGF thereby creating a pathological positive feedback loop that continues to increase integrins^35, 68^. In addition, increased VEGFR2-PI3K signaling promotes activation of integrins^35, 64^, and integrins in focal adhesions activate YAP/TAZ through Rho GTPase (CDC42) activation^32^. Thus, ALK1 is an important regulator of VEGFR2-PI3K-integrin-YAP/TAZ positive feedback loop. While mechanistic details of pathway interaction remain to be determined, our data identify the YAP/TAZ inhibitor Verteporfin, which inhibits YAP/TAZ translocation to the nucleus to block YAP-TEAD association^65, 66^, as a novel inhibitor therapy for AVMs. Verteporfin photodynamic therapy is approved for the treatment of choroidal neovascularization due to age-related macular degeneration^67^ and both integrin inhibitors and verteporfin might be novel therapeutic options for HHT patients.

## Acknowledgements

We thank ATTRACT members Paul Oh, Lena Claesson-Welsh, Miguel Bernabeu and Holger Gerhardt for critical comments on the manuscript, Profs Bin Zhou (Shanghai Institute for Biological Sciences) and Ralf Adams (Max-Planck Institute, Munster, Germany) for mouse lines, and Dr. Jihoon Park for Python scripts to measure polarity and index.

## Affiliations

Cardiovascular Research Center, Department of Internal Medicine, Yale University School of Medicine, New Haven CT, USA. (H.P, J.F, M.P, M.C, S.Y, M.A.S, A.E), Yale University School of Medicine, Department of Pharmacology (L.S, W.C.S), Instituto de Medicina Molecular João Lobo Antunes, Faculdade de Medicina, Universidade de Lisboa, Lisboa, Portugal (C.F), Yale University School of Medicine, Departments of Cell Biology and Biomedical Engineering (M.A.S), Yale University School of Medicine, Department of Molecular and Cellular Physiology (A.E), Université de Paris, PARCC, INSERM, F-75006 Paris (A.E)

---

**Supplemental Figure 1.**
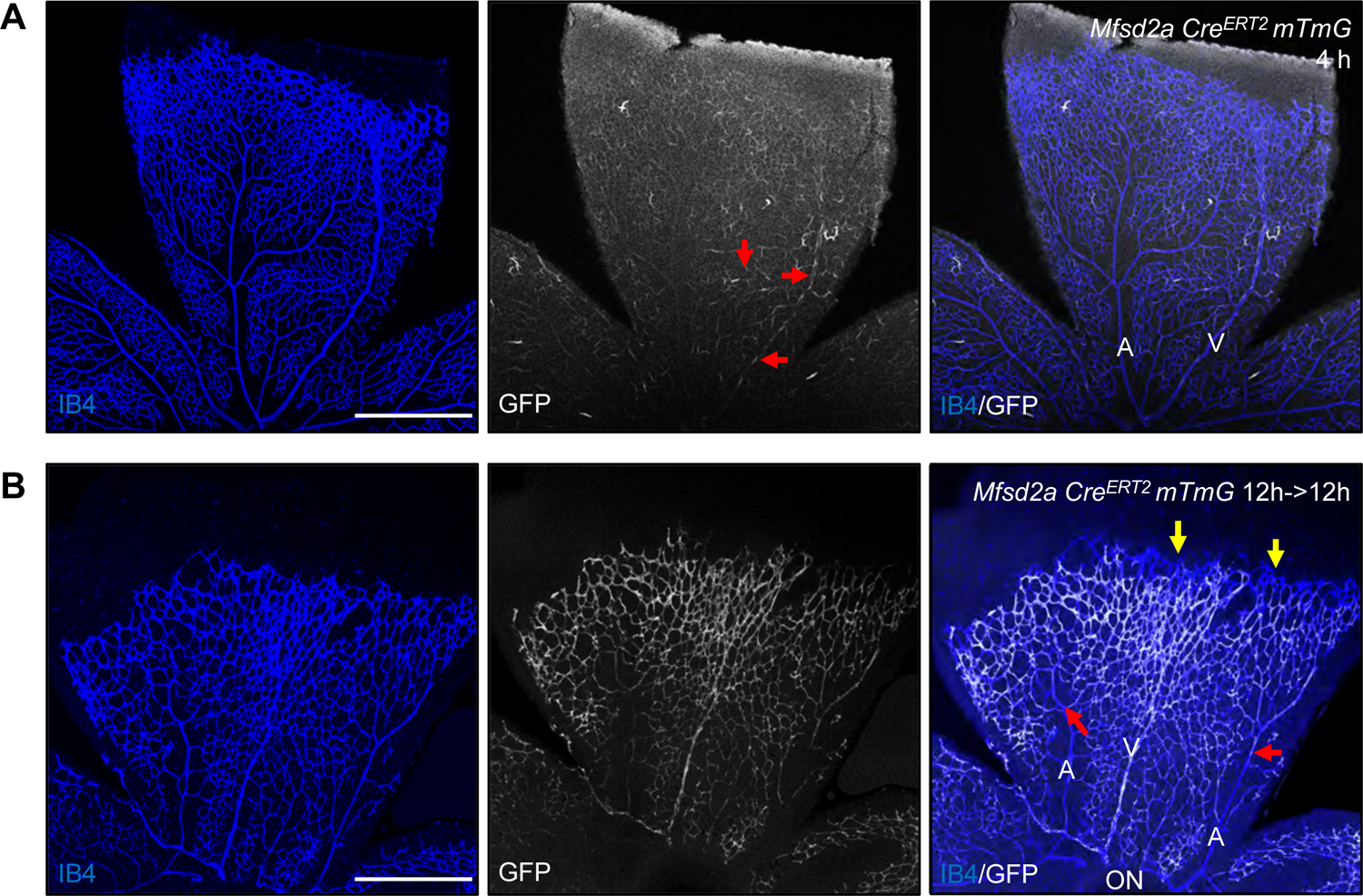
*Mfsd2a* positive cells migrate against the direction of blood flow. (A) *Mfsd2a Cre^ERT2^ mTmG* mice injected with 100 μg Tx at P6 and dissected after 4 h. GFP expressing ECs are located in capillaries and veins (red arrows) but not in arteries. (B) P6 retina flat mount images labeled with IB4 (blue) and GFP (white) from *Mfsd2a Cre^ERT2^ mTmG* mice injected with 100 μg Tx at P5, dissected after 12h (P5.5) and cultured for an additional 12 h in vitro (P6). Yellow arrows indicate tip cells and red arrows indicate location of GFP-expressing ECs in arteries. ON: optic nerve, V: vein, A: artery, Scale bar: 500 μm

**Supplemental Figure 2.**
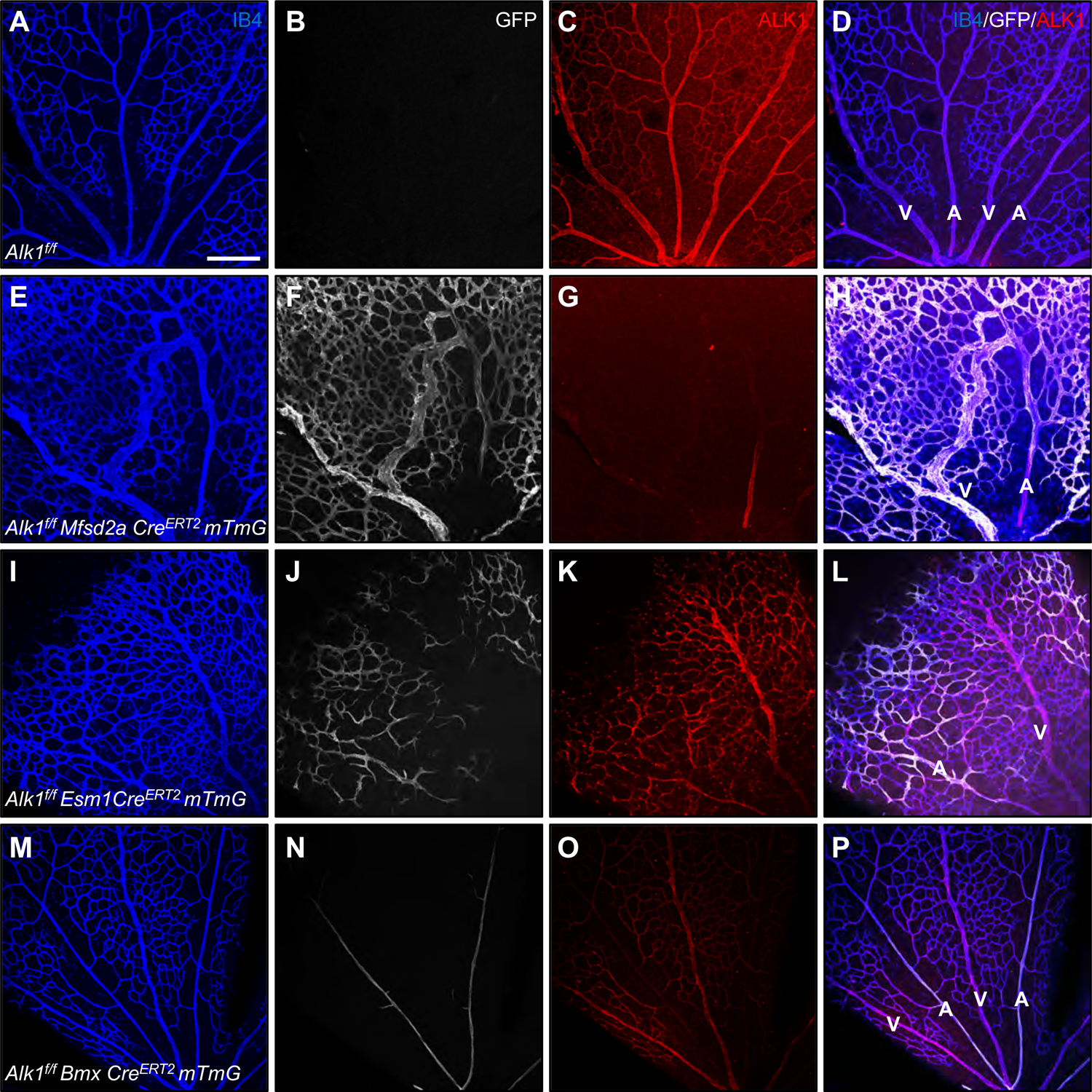
Genetic deletion of *Alk1* in *Mfsd2a*, *Esm1* and *Bmx Cre^ERT2^ mTmG* retinas. (A-P) 100 μg Tx was injected intragastrically at P4 in *Alk1^f/f^*, *Alk1^f/f^ Mfsd2a Cre^ERT2^*, *Alk1^f/f^ Esm1 Cre^ERT2^* and *Alk1^f/f^ Bmx Cre^ERT2^ mTmG* pups, and retinas were dissected at P6. IB4 (blue), GFP (white) and ALK1 (red) staining of retinal flat mounts. GFP and ALK1 staining shows non-overlapping expression. V: vein, A: artery, Scale bar: 200 μm

**Supplemental Figure 3.**
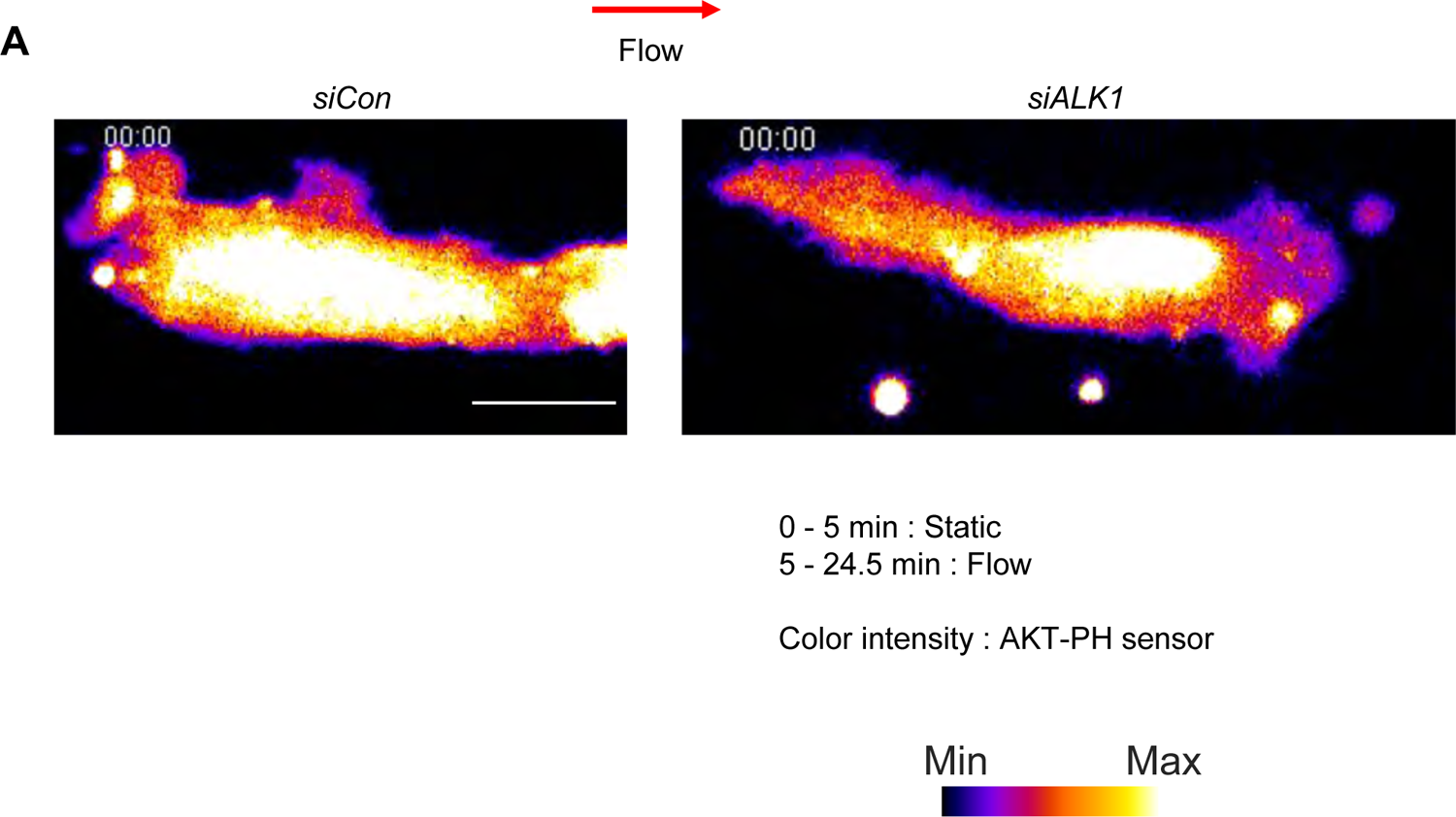
ALK1 regulates EC polarization against the direction of blood flow. Movies of *siCon* or *siALK1* HUVECs stably transduced with PH-AKT-mClover3 and plasma membrane targeting sequence of LCK-mRuby3. HUVEC monolayers in microfluidic chambers were exposed to 12 dynes/cm^2^ LSS under microscope. 0 – 5 min is static condition and flow starts after 5 min. Color intensity indicates AKT-PH sensor. A scale bar: 20 μm

**Supplemental Figure 4.**
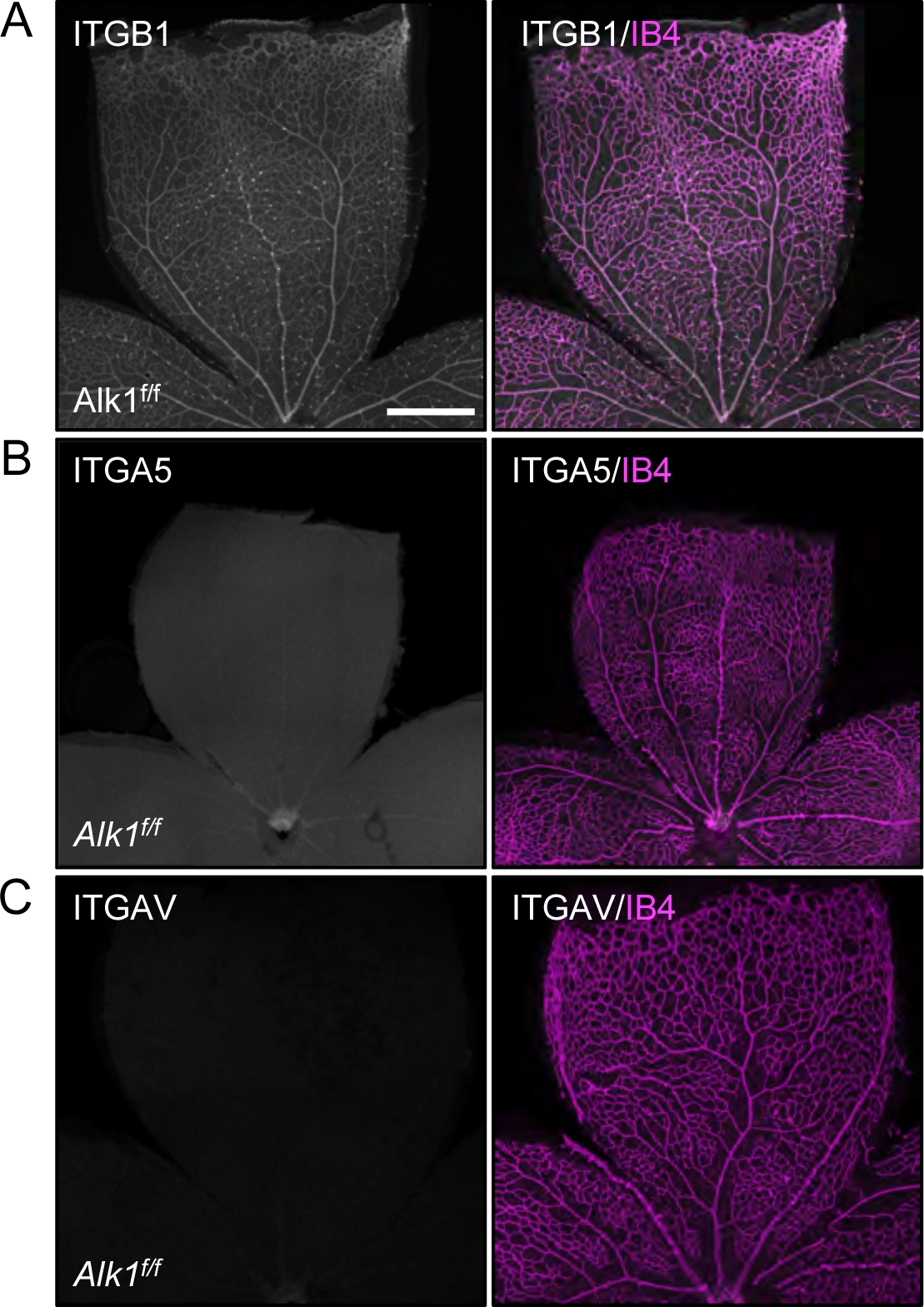
Integrin staining of *Alk1^f/f^* retinas. (A-C) IB4 (Magenta) and ITGB1(A, white), ITGA5 (B, white) or ITGAV (C, white) staining of retinal flat mounts from P8 *Alk1^f/f^* pups. A scale bar: 500 μm (A-C)

**Supplemental Figure 5.**
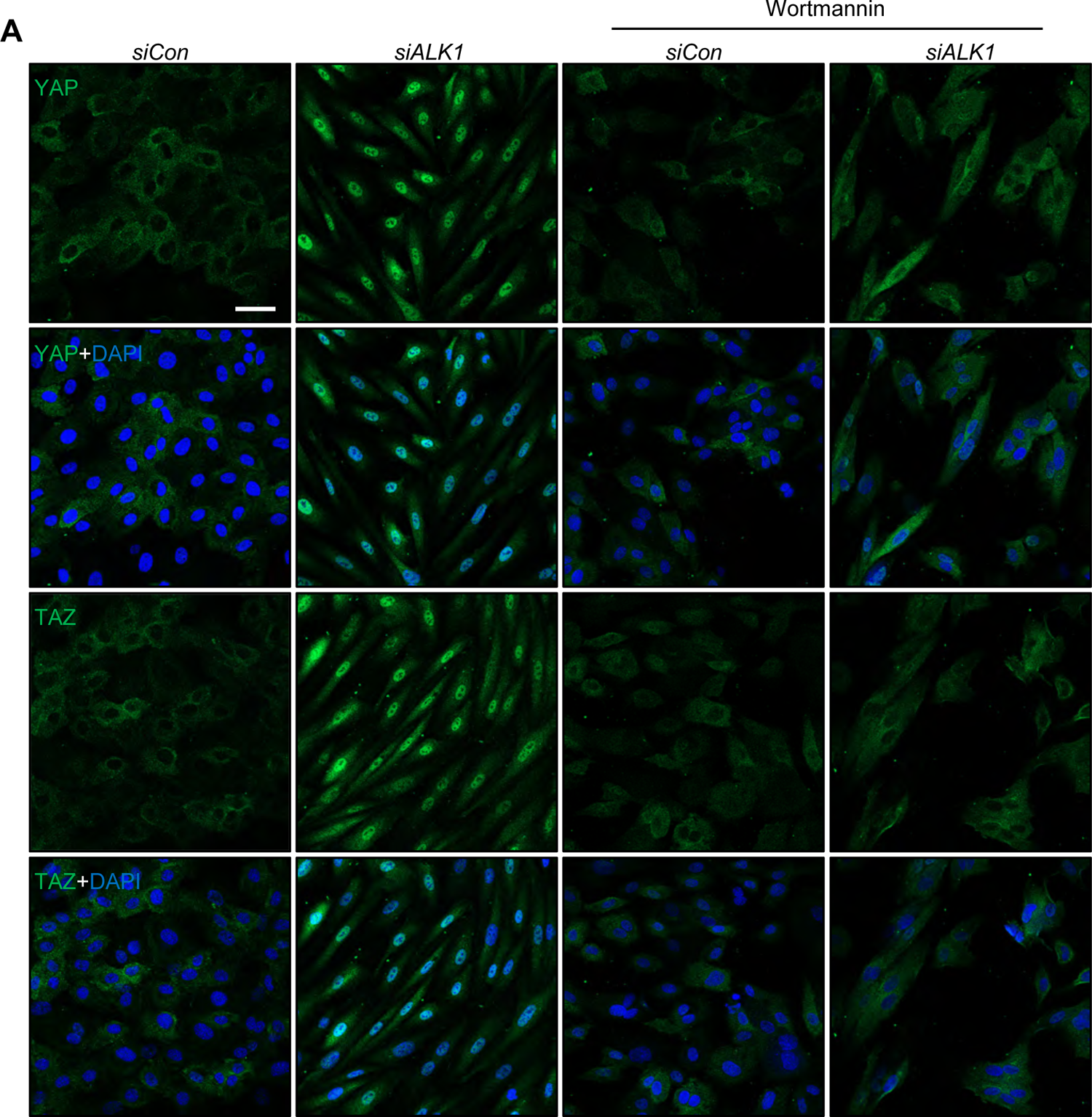
PI3K acts upstream of YAP/TAZ. (A) YAP and TAZ staining for control, *ALK1* siRNAs transfected HUVECs treated with PBS, Wortmannin (100 nM) for 12 h. A scale bar: 50 μm

## References

1. Shovlin CL. Hereditary haemorrhagic telangiectasia: Pathophysiology, diagnosis and treatment. Blood Reviews. 2010;24:203–219.

2. McAllister KA, Grogg KM, Johnson DW, Gallione CJ, Baldwin MA, Jackson CE, Helmbold EA, Markel DS, McKinnon WC, Murrel J, McCormick MK, Pericak-Vance MA, Heutink P, Oostra BA, Haitjema T, Westerman CJJ, Porteous ME, Guttmacher AE, Letarte M and Marchuk DA. Endoglin, a TGF-β binding protein of endothelial cells, is the gene for hereditary haemorrhagic telangiectasia type 1. Nature Genetics. 1994;8:345–351.

3. Johnson DW, Berg JN, Baldwin MA, Gallione CJ, Marondel I, Yoon SJ, Stenzel TT, Speer M, Pericak-Vance MA, Diamond A, Guttmacher AE, Jackson CE, Attisano L, Kucherlapati R, Porteous MEM and Marchuk DA. Mutations in the activin receptor–like kinase 1 gene in hereditary haemorrhagic telangiectasia type 2. Nature Genetics. 1996;13:189–195.

4. Gallione CJ, Repetto GM, Legius E, Rustgi AK, Schelley SL, Tejpar S, Mitchell G, Drouin E, Westermann CJ and Marchuk DA. A combined syndrome of juvenile polyposis and hereditary haemorrhagic telangiectasia associated with mutations in MADH4 (SMAD4). Lancet. 2004;363:852–9.

5. Gallione CJ, Klaus DJ, Yeh EY, Stenzel TT, Xue Y, Anthony KB, McAllister KA, Baldwin MA, Berg JN, Lux A, Smith JD, Vary CPH, Craigen WJ, Westermann CJJ, Warner ML, Miller YE, Jackson CE, Guttmacher AE and Marchuk DA. Mutation and expression analysis of the endoglin gene in Hereditary Hemorrhagic Telangiectasia reveals null alleles. Human Mutation. 1998;11:286–294.

6. Roman BL and Hinck AP. ALK1 signaling in development and disease: new paradigms. Cellular and Molecular Life Sciences. 2017;74:4539–4560.

7. David L, Mallet C, Mazerbourg S, Feige JJ and Bailly S. Identification of BMP9 and BMP10 as functional activators of the orphan activin receptor-like kinase 1 (ALK1) in endothelial cells. Blood. 2007;109:1953–61.

8. Ruiz-Llorente L, Gallardo-Vara E, Rossi E, Smadja DM, Botella LM and Bernabeu C. Endoglin and alk1 as therapeutic targets for hereditary hemorrhagic telangiectasia. Expert Opinion on Therapeutic Targets. 2017;21:933–947.

9. Snellings DA, Gallione CJ, Clark DS, Vozoris NT, Faughnan ME and Marchuk DA. Somatic Mutations in Vascular Malformations of Hereditary Hemorrhagic Telangiectasia Result in Bi-allelic Loss of ENG or ACVRL1. American Journal of Human Genetics. 2019;105:894–906.

10. Govani FS and Shovlin CL. Hereditary haemorrhagic telangiectasia: A clinical and scientific review. European Journal of Human Genetics. 2009;17:860–871.

11. McDonald J, Bayrak-Toydemir P and Pyeritz RE. Hereditary hemorrhagic telangiectasia: An overview of diagnosis, management, and pathogenesis. Genetics in Medicine. 2011;13:607–616.

12. Tual-Chalot S, Mahmoud M, Allinson KR, Redgrave RE, Zhai Z, Oh SP, Fruttiger M and Arthur HM. Endothelial depletion of Acvrl1 in mice leads to arteriovenous malformations associated with reduced endoglin expression. PLoS One. 2014;9:e98646.

13. Ola R, Dubrac A, Han J, Zhang F, Fang JS, Larrivée B, Lee M, Urarte AA, Kraehling JR, Genet G, Hirschi KK, Sessa WC, Canals FV, Graupera M, Yan M, Young LH, Oh PS and Eichmann A. PI3 kinase inhibition improves vascular malformations in mouse models of hereditary haemorrhagic telangiectasia. Nature Communications. 2016;7:13650.

14. Ola R, Künzel Sandrine H, Zhang F, Genet G, Chakraborty R, Pibouin-Fragner L, Martin K, Sessa W, Dubrac A and Eichmann A. SMAD4 Prevents Flow Induced Arteriovenous Malformations by Inhibiting Casein Kinase 2. Circulation. 2018;138:2379–2394.

15. Baeyens N, Larrivée B, Ola R, Hayward-Piatkowskyi B, Dubrac A, Huang B, Ross TD, Coon BG, Min E, Tsarfati M, Tong H, Eichmann A and Schwartz MA. Defective fluid shear stress mechanotransduction mediates hereditary hemorrhagic telangiectasia. The Journal of Cell Biology. 2016;214:807.

16. Capasso TL, Li B, Volek HJ, Khalid W, Rochon ER, Anbalagan A, Herdman C, Yost HJ, Villanueva FS, Kim K and Roman BL. BMP10-mediated ALK1 signaling is continuously required for vascular development and maintenance. Angiogenesis. 2020;23:203–220.

17. Rochon ER, Menon PG and Roman BL. Alk1 controls arterial endothelial cell migration in lumenized vessels. Development. 2016;143:2593–602.

18. Han C, Choe S-w, Kim YH, Acharya AP, Keselowsky BG, Sorg BS, Lee Y-J and Oh SP. VEGF neutralization can prevent and normalize arteriovenous malformations in an animal model for hereditary hemorrhagic telangiectasia 2. Angiogenesis. 2014;17:823–830.

19. Jin Y, Muhl L, Burmakin M, Wang Y, Duchez AC, Betsholtz C, Arthur HM and Jakobsson L. Endoglin prevents vascular malformation by regulating flow-induced cell migration and specification through VEGFR2 signalling. Nature Cell Biology. 2017;19:639–652.

20. Alsina-Sanchís E, García-Ibáñez Y, Figueiredo AM, Riera-Domingo C, Figueras A, Matias-Guiu X, Casanovas O, Botella LM, Pujana MA, Riera-Mestre A, Graupera M and Viñals F. ALK1 Loss Results in Vascular Hyperplasia in Mice and Humans Through PI3K Activation. Arterioscler Thromb Vasc Biol. 2018;38:1216–1229.

21. Iriarte A, Figueras A, Cerdà P, Mora JM, Jucglà A, Penín R, Viñals F and Riera-Mestre A. PI3K (Phosphatidylinositol 3-Kinase) Activation and Endothelial Cell Proliferation in Patients with Hemorrhagic Hereditary Telangiectasia Type 1. Cells. 2019;8.

22. Carvalho JR, Fortunato IC, Fonseca CG, Pezzarossa A, Barbacena P, Dominguez-Cejudo MA, Vasconcelos FF, Santos NC, Carvalho FA and Franco CA. Non-canonical Wnt signaling regulates junctional mechanocoupling during angiogenic collective cell migration. eLife. 2019;8:e45853.

23. Pu W, Zhang H, Huang X, Tian X, He L, Wang Y, Zhang L, Liu Q, Li Y, Li Y, Zhao H, Liu K, Lu J, Zhou Y, Huang P, Nie Y, Yan Y, Hui L, Lui KO and Zhou B. Mfsd2a+ hepatocytes repopulate the liver during injury and regeneration. Nat Commun. 2016;7:13369.

24. Pu W, He L, Han X, Tian X, Li Y, Zhang H, Liu Q, Huang X, Zhang L, Wang QD, Yu Z, Yang X, Smart N and Zhou B. Genetic Targeting of Organ-Specific Blood Vessels. Circ Res. 2018;123:86–99.

25. Chow BW, Nuñez V, Kaplan L, Granger AJ, Bistrong K, Zucker HL, Kumar P, Sabatini BL and Gu C. Caveolae in CNS arterioles mediate neurovascular coupling. Nature. 2020;579:106–110.

26. Pitulescu ME, Schmidt I, Giaimo BD, Antoine T, Berkenfeld F, Ferrante F, Park H, Ehling M, Biljes D, Rocha SF, Langen UH, Stehling M, Nagasawa T, Ferrara N, Borggrefe T and Adams RH. Dll4 and Notch signalling couples sprouting angiogenesis and artery formation. Nature Cell Biology. 2017;19:915.

27. Xu C, Hasan SS, Schmidt I, Rocha SF, Pitulescu ME, Bussmann J, Meyen D, Raz E, Adams RH and Siekmann AF. Arteries are formed by vein-derived endothelial tip cells. Nature Communications. 2014;5:5758.

28. Ehling M, Adams S, Benedito R and Adams RH. Notch controls retinal blood vessel maturation and quiescence. Development. 2013;140:3051.

29. Muzumdar MD, Tasic B, Miyamichi K, Li L and Luo L. A global double-fluorescent Cre reporter mouse. Genesis. 2007;45:593–605.

30. Melchior B and Frangos JA. Distinctive subcellular Akt-1 responses to shear stress in endothelial cells. Journal of Cellular Biochemistry. 2014;115:121–129.

31. Várnai P and Balla T. Visualization of phosphoinositides that bind pleckstrin homology domains: Calcium- and agonist-induced dynamic changes and relationship to myo-[3H]inositol-labeled phosphoinositide pools. Journal of Cell Biology. 1998;143:501–510.

32. Etienne-Manneville S and Hall A. Integrin-mediated activation of Cdc42 controls cell polarity in migrating astrocytes through PKCzeta. Cell. 2001;106:489–98.

33. Tzima E, Kiosses WB, del Pozo MA and Schwartz MA. Localized cdc42 activation, detected using a novel assay, mediates microtubule organizing center positioning in endothelial cells in response to fluid shear stress. J Biol Chem. 2003;278:31020-3.

34. Tzima E, Irani-Tehrani M, Kiosses WB, Dejana E, Schultz DA, Engelhardt B, Cao G, DeLisser H and Schwartz MA. A mechanosensory complex that mediates the endothelial cell response to fluid shear stress. Nature. 2005;437:426–31.

35. Somanath PR, Malinin NL and Byzova TV. Cooperation between integrin ανβ3 and VEGFR2 in angiogenesis. Angiogenesis. 2009;12:177–185.

36. Simons M. An inside view: VEGF receptor trafficking and signaling. Physiology (Bethesda). 2012;27:213–22.

37. Serini G, Napione L, Arese M and Bussolino F. Besides adhesion: new perspectives of integrin functions in angiogenesis. Cardiovascular Research. 2008;78:213–222.

38. Neto F, Klaus-Bergmann A, Ong YT, Alt S, Vion A-C, Szymborska A, Carvalho JR, Hollfinger I, Bartels-Klein E, Franco CA, Potente M and Gerhardt H. YAP and TAZ regulate adherens junction dynamics and endothelial cell distribution during vascular development. eLife. 2018;7:e31037.

39. Nakajima H, Yamamoto K, Agarwala S, Terai K, Fukui H, Fukuhara S, Ando K, Miyazaki T, Yokota Y, Schmelzer E, Belting H-G, Affolter M, Lecaudey V and Mochizuki N. Flow-Dependent Endothelial YAP Regulation Contributes to Vessel Maintenance. Developmental Cell. 2017;40:523–536.e6.

40. Wang L, Luo JY, Li B, Tian XY, Chen LJ, Huang Y, Liu J, Deng D, Lau CW, Wan S, Ai D, Mak KK, Tong KK, Kwan KM, Wang N, Chiu JJ, Zhu Y and Huang Y. Integrin-YAP/TAZ-JNK cascade mediates atheroprotective effect of unidirectional shear flow. Nature. 2016;540:579–582.

41. Li B, He J, Lv H, Liu Y, Lv X, Zhang C, Zhu Y and Ai D. c-Abl regulates YAPY357 phosphorylation to activate endothelial atherogenic responses to disturbed flow. J Clin Invest. 2019;129:1167–1179.

42. Dupont S. Role of YAP/TAZ in cell-matrix adhesion-mediated signalling and mechanotransduction. Exp Cell Res. 2016;343:42–53.

43. Sakabe M, Fan J, Odaka Y, Liu N, Hassan A, Duan X, Stump P, Byerly L, Donaldson M, Hao J, Fruttiger M, Lu QR, Zheng Y, Lang RA and Xin M. YAP/TAZ-CDC42 signaling regulates vascular tip cell migration. Proc Natl Acad Sci U S A. 2017;114:10918–10923.

44. Kim J, Kim YH, Kim J, Park DY, Bae H, Lee DH, Kim KH, Hong SP, Jang SP, Kubota Y, Kwon YG, Lim DS and Koh GY. YAP/TAZ regulates sprouting angiogenesis and vascular barrier maturation. J Clin Invest. 2017;127:3441–3461.

45. Franco CA, Jones ML, Bernabeu MO, Geudens I, Mathivet T, Rosa A, Lopes FM, Lima AP, Ragab A, Collins RT, Phng L-K, Coveney PV and Gerhardt H. Dynamic Endothelial Cell Rearrangements Drive Developmental Vessel Regression. PLOS Biology. 2015;13:e1002125.

46. Fonseca CG, Barbacena P and Franco CA. Endothelial cells on the move: dynamics in vascular morphogenesis and disease. Vasc Biol. 2020;2:H29–h43.

47. Chow BW and Gu C. Gradual Suppression of Transcytosis Governs Functional Blood-Retinal Barrier Formation. Neuron. 2017;93:1325–1333.e3.

48. Singh E, Redgrave RE, Phillips HM and Arthur HM. Arterial endoglin does not protect against arteriovenous malformations. Angiogenesis. 2020;23:559–566.

49. Corti P, Young S, Chen CY, Patrick MJ, Rochon ER, Pekkan K and Roman BL. Interaction between alk1 and blood flow in the development of arteriovenous malformations. Development. 2011;138:1573–82.

50. Tual-Chalot S, Garcia-Collado M, Redgrave RE, Singh E, Davison B, Park C, Lin H, Luli S, Jin Y, Wang Y, Lawrie A, Jakobsson L and Arthur HM. Loss of endothelial endoglin promotes high-output heart failure through peripheral arteriovenous shunting driven by VEGF signaling. Circulation Research. 2020:243–257.

51. Thalgott JH, Dos-Santos-Luis D, Hosman AE, Martin S, Lamandé N, Bracquart D, Srun S, Galaris G, De Boer HC, Tual-Chalot S, Kroon S, Arthur HM, Cao Y, Snijder RJ, Disch F, Mager JJ, Rabelink TJ, Mummery CL, Raymond K and Lebrin F. Decreased expression of vascular endothelial growth factor receptor 1 contributes to the pathogenesis of hereditary hemorrhagic telangiectasia type 2. Circulation. 2018;138:2698–2712.

52. Ruiz S, Zhao H, Chandakkar P, Papoin J, Choi H, Nomura-Kitabayashi A, Patel R, Gillen M, Diao L, Chatterjee PK, He M, Al-Abed Y, Wang P, Metz CN, Oh SP, Blanc L, Campagne F and Marambaud P. Correcting Smad1/5/8, mTOR, and VEGFR2 treats pathology in hereditary hemorrhagic telangiectasia models. Journal of Clinical Investigation. 2020;130:942-957.

53. Hwan Kim Y, Vu PN, Choe SW, Jeon CJ, Arthur HM, Vary CPH, Lee YJ and Oh SP. Overexpression of Activin Receptor-Like Kinase 1 in Endothelial Cells Suppresses Development of Arteriovenous Malformations in Mouse Models of Hereditary Hemorrhagic Telangiectasia. Circ Res. 2020;127:1122–1137.

54. Crist AM, Zhou X, Garai J, Lee AR, Thoele J, Ullmer C, Klein C, Zabaleta J and Meadows SM. Angiopoietin-2 Inhibition Rescues Arteriovenous Malformation in a Smad4 Hereditary Hemorrhagic Telangiectasia Mouse Model. Circulation. 2019;139:2049–2063.

55. Xanthis I, Souilhol C, Serbanovic-Canic J, Roddie H, Kalli AC, Fragiadaki M, Wong R, Shah DR, Askari JA, Canham L, Akhtar N, Feng S, Ridger V, Waltho J, Pinteaux E, Humphries MJ, Bryan MT and Evans PC. β1 integrin is a sensor of blood flow direction. Journal of Cell Science. 2019;132.

56. Urbich C, Walter DH, Zeiher AM and Dimmeler S. Laminar shear stress upregulates integrin expression role in endothelial cell adhesion and apoptosis. Circulation Research. 2000;87:683–689.

57. Tzima E, Del Pozo MA, Shattil SJ, Chien S and Schwartz MA. Activation of integrins in endothelial cells by fluid shear stress mediates Rho-dependent cytoskeletal alignment. EMBO Journal. 2001;20:4639–4647.

58. Stupp R, Hegi ME, Gorlia T, Erridge SC, Perry J, Hong YK, Aldape KD, Lhermitte B, Pietsch T, Grujicic D, Steinbach JP, Wick W, Tarnawski R, Nam DH, Hau P, Weyerbrock A, Taphoorn MJ, Shen CC, Rao N, Thurzo L, Herrlinger U, Gupta T, Kortmann RD, Adamska K, McBain C, Brandes AA, Tonn JC, Schnell O, Wiegel T, Kim CY, Nabors LB, Reardon DA, van den Bent MJ, Hicking C, Markivskyy A, Picard M, Weller M, European Organisation for R, Treatment of C, Canadian Brain Tumor C and team Cs. Cilengitide combined with standard treatment for patients with newly diagnosed glioblastoma with methylated MGMT promoter (CENTRIC EORTC 26071-22072 study): a multicentre, randomised, open-label, phase 3 trial. The Lancet Oncology. 2014;15:1100-1108.

59. Wang X, Freire Valls A, Schermann G, Shen Y, Moya IM, Castro L, Urban S, Solecki GM, Winkler F, Riedemann L, Jain RK, Mazzone M, Schmidt T, Fischer T, Halder G and Ruiz de Almodóvar C. YAP/TAZ Orchestrate VEGF Signaling during Developmental Angiogenesis. Developmental Cell. 2017;42:462–478.e7.

60. Wang K-C, Yeh Y-T, Nguyen P, Limqueco E, Lopez J, Thorossian S, Guan K-L, Li Y-SJ and Chien S. Flow-dependent YAP/TAZ activities regulate endothelial phenotypes and atherosclerosis. Proceedings of the National Academy of Sciences. 2016;113:11525.

61. Kim J, Kim YH, Kim J, Park DY, Bae H, Lee D-H, Kim KH, Hong SP, Jang SP, Kubota Y, Kwon Y-G, Lim D-S and Koh GY. YAP/TAZ regulates sprouting angiogenesis and vascular barrier maturation. The Journal of Clinical Investigation. 2017;127:3441–3461.

62. Boopathy GTK and Hong W. Role of Hippo Pathway-YAP/TAZ signaling in angiogenesis. Frontiers in Cell and Developmental Biology. 2019;7.

63. Laviña B, Castro M, Niaudet C, Cruys B, Álvarez-Aznar A, Carmeliet P, Bentley K, Brakebusch C, Betsholtz C and Gaengel K. Defective endothelial cell migration in the absence of Cdc42 leads to capillary-venous malformations. Development (Cambridge). 2018;145.

64. Myo-Hyeon P, Kim AK, Manandhar S, Su-Young O, Gun-Hyuk J, Kang L, Dong-Won L, Hyeon DY, Sun-Hee L, Lee HE, Tae-Lin H, Heon Suh S, Hwang D, Byun K, Hae-Chul P and Lee YM. CCN1 interlinks integrin and hippo pathway to autoregulate tip cell activity. eLife. 2019;8.

65. Gibault F, Bailly F, Corvaisier M, Coevoet M, Huet G, Melnyk P and Cotelle P. Molecular Features of the YAP Inhibitor Verteporfin: Synthesis of Hexasubstituted Dipyrrins as Potential Inhibitors of YAP/TAZ, the Downstream Effectors of the Hippo Pathway. ChemMedChem. 2017;12:954–961.

66. Chan WM, Lim TH, Pece A, Silva R and Yoshimura N. Verteporfin PDT for non-standard indications-a review of current literature. Graefe’s Archive for Clinical and Experimental Ophthalmology. 2010;248:613–626.

67. Messmer KJ and Abel SR. Verteporfin for Age-Related Macular Degeneration. Annals of Pharmacotherapy. 2001;35:1593–1598.

68. Kim N-G and Gumbiner BM. Adhesion to fibronectin regulates Hippo signaling via the FAK–Src–PI3K pathway. The Journal of Cell Biology. 2015;210:503.

